# A nanoscale reciprocating rotary mechanism with coordinated mobility control

**DOI:** 10.1101/2021.04.27.441587

**Authors:** Eva Bertosin, Christopher M. Maffeo, Thomas Drexler, Maximilian N. Honemann, Aleksei Aksimentiev, Hendrik Dietz

## Abstract

Biological molecular motors transform chemical energy into mechanical work by coupling a cycle of catalytic reactions to large scale structural transitions. Mechanical deformation can be surprisingly efficient in realizing such coupling, as demonstrated by the celebrated example of F_1_F_0_ ATP synthase. Here, we describe a synthetic molecular mechanism that transforms a rotary motion of an asymmetric camshaft into reciprocating large-scale transitions in the structure of the surrounding stator orchestrated by mechanical deformation. We designed the mechanism using DNA origami, characterized the structure of the components and the entire mechanism using cryo-electron microscopy, and examined the mechanism’s dynamic behavior using single-particle fluorescence microscopy and molecular dynamics simulations. The data indicates that, while the camshaft can rotate inside the stator by diffusion, mechanical deformation of the stator makes the camshaft pause at a set of preferred orientations. By changing the mechanical stiffness of the stator, we could accelerate or suppress the Brownian rotation within the mechanism, thereby demonstrating an allosteric coupling between the movement of the camshaft and of the stator, and the ability to tailor the free energy landscape that governs the rotary motion. Our mechanism provides a framework for the manufacture of artificial nanomachines that, just like the man-made machines in the macroscopic world, function because of coordinated movements of their components.

## Introduction

Macroscopic machines commonly rely on a coordinated motion of multiple rigid components to perform their function. For example, an internal combustion engine uses a rotating camshaft to cyclically open or close the peripheral valves for fuel injection and exhaust gas removal; the coordination of the valves’ operation is paramount to the engine’s function. Nanoscale biological machines also often consist of multiple components that move in a coordinated fashion. For example, the rotation of the central shaft in F_1_F_0_ ATP synthase (*1-3*) produces cyclic structural transformations at the interfaces of the F_1_ subunits, coordinating cyclic chemical transformations. Intriguingly, the F_1_ motor of F_1_F_0_ ATP synthase is fully reversible (*4, 5*): it can either function as a rotary motor powered by the chemical energy of ATP hydrolysis or it can use the mechanical energy of the central shaft rotation to catalyze synthesis of ATP. The fact that F_1_ ATP synthase can be both a motor and a chemical generator reflects the microscopic reversibility of elementary chemical processes and is a unique feature of molecular scale machines. Realizing a similar degree of mechanochemical coupling in a synthetic nanoscale system remains a landmark technological goal.

The construction of artificial molecular machines by chemical synthesis has previously provided important insights regarding how to create molecular mechanisms with internal degrees of freedom, such as catenanes and rotaxanes, and how to power molecular motions using chemical fuels, light, and other stimuli (*6-12*). DNA nanotechnology has also already provided a range of mechanical systems including pivots, hinges, crank sliders, and rotary mechanisms (*13-16*) that can be reconfigured using strand displacement reactions (SDR) (*17*) or by changing environmental parameters such as pH, ionic strength, temperature, and external fields (*18-23*). Whereas the molecular mechanisms generated by chemical synthesis tend to include on the order of 100 atoms, DNA nanostructures, in particular DNA origami objects, are much larger and can encompass hundreds of thousands of atoms (*24-26*). As such, DNA origami nanomachines may offer more opportunities for the assembly of space-filling components that realize mechanisms of coordinated mechanical power transmission. In this work, we describe the construction, computational characterization and experimental validation of a rotary mechanism with user-defined power and motion transmission. We conceived this object by combining macroscale machine design concepts with functional and structural aspects of the ATP synthase, and consider it to be an important milestone toward creating artificial machines that achieve and generalize functionalities observed in biological motors.

## Results

### Design of the rotary mechanism

We designed our rotary mechanism as a tetramer composed of a camshaft-like rotor in a surrounding stator. Rotations of the camshaft will induce reciprocal deformations of the structural elements in the stator (Fig. 1A). The stator is made from three similarly shaped components. These stator units are designed to contain rigid parts forming a “bearing” which will hold the shaft. Each unit of the stator also has two “pawls” that can flex in response to the camshaft rotation (Fig. 1B). In an implementation where each of the pawls can flex independently, the full stator unit features six preferred camshaft orientation slots. As we will discuss, the pawls can also be coupled together to reduce the number of available slots, which will influence the motion of the camshaft. For example, Fig. 1A shows the three available slots when the two pawls per stator unit were to flex together. The camshaft also contains a crossbar (Fig. 1C). The crossbar and the cam on the other end of the shaft are used to mechanically interlock the camshaft inside the stator (Fig. 1D).

**Figure 1.**
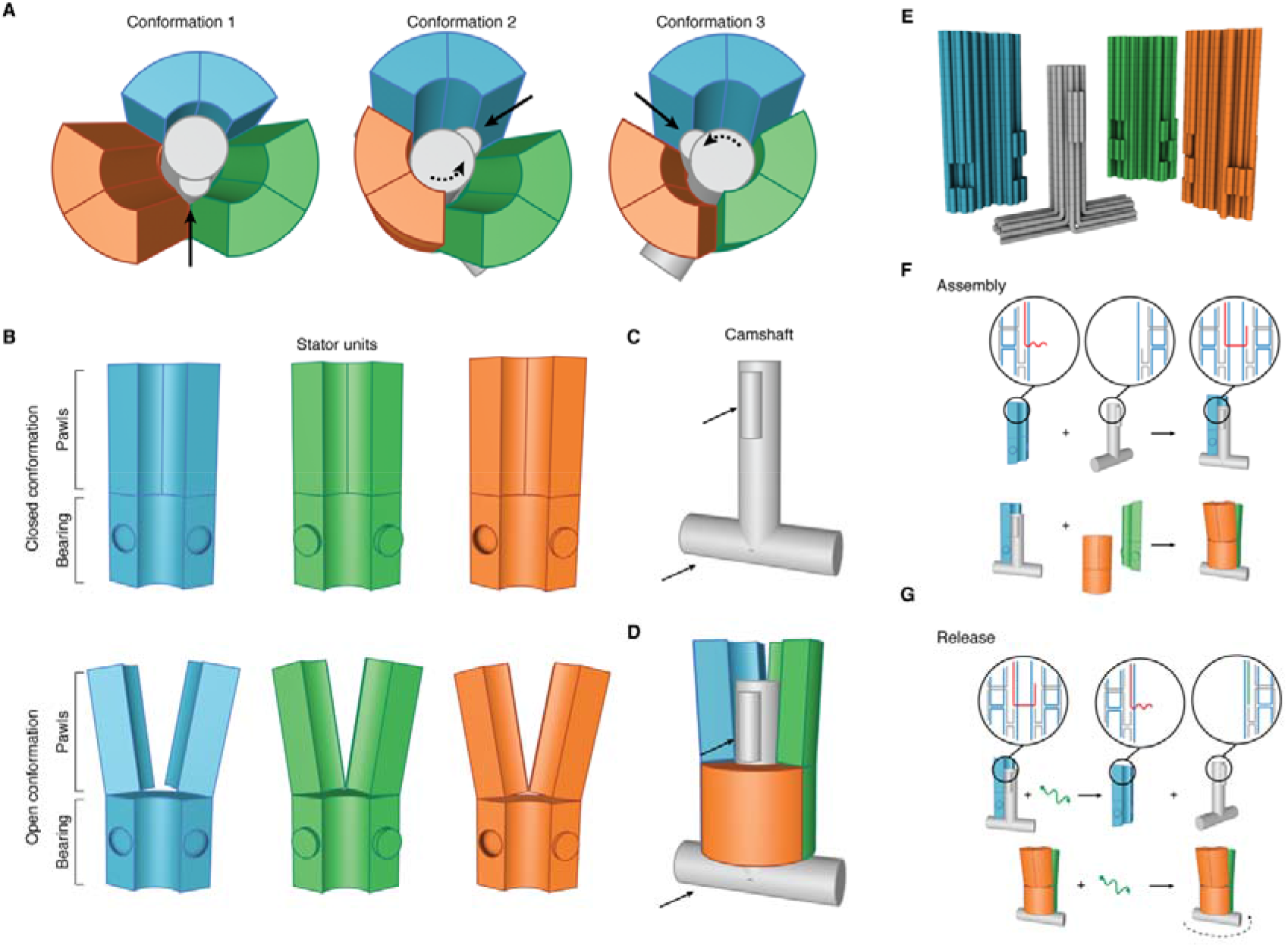
Conceptual design and assembly of the rotary mechanism. **(A)** Sketches of the mechanism. The shaft is depicted in gray, while the stator units are in blue, green and orange. **(B)** Sketches of the stator units with pawls in the closed (top) and open (bottom) conformations. **(C)** Sketch of the camshaft. Black arrows indicate features used to prevent the camshaft from escaping the stator. **(D)** Sketch of the camshaft when trapped inside the stator. Pawls of the orange stator unit are not drawn. Black arrows as in (C). **(E)** Cylinder models of our implementation with multilayer DNA origami. Cylinders represent DNA double helices. **(F)** Sketch of the assembly steps for building the rotary mechanism. Red: connecting strands. **(G)** Top: sketch of the shaft release from the stator unit via toehold-mediated strand displacement. Bottom: same reaction but performed within the fully assembled stator. Green: invader strands.

We approximated the desired 3D shapes of the components using the methods of multilayer DNA origami (Fig. 1E) (*27, 28*). In our design, each stator unit consists of 46 helices packed in parallel on a honeycomb-like lattice. One stator unit possesses an asymmetric feature to allow discriminating the stator orientation relative to the shaft orientation by transmission electron microscopy (TEM) imaging (Fig. S1). The bearing and the pawls can be considered each as rigid blocks of helices that are connected to each other via two DNA double helices (the “support” helices) that extend vertically along the whole structure. The pawls are connected to the support helices via two crossovers at the top and bottom of the pawls, respectively. The support helices can flex away from the central shaft, and the pawls can also flex around the support helices. This construction offers two elastic degrees of freedom to make room for the rotating camshaft. The pawls can also form base-pair stacking interactions at the blunt-ended helical interfaces between the bottom and top surface of pawls and bearing, respectively. These interactions can be utilized to influence the flexibility of the pawls.

The camshaft consists of a shaft made of 24 helices packed in a honeycomb-like lattice. The T-crossbar is formed by helices that are bent to form a 90° angle with the shaft. The cam on the shaft is produced by four additional helices that are attached on one side of the shaft (Fig. 1E, Fig. S2). The cross section of the shaft fits tightly into the central bore of the bearing (Fig. S3), but the shaft’s protruding feature clashes with the pawl helices, forcing them to flex away from the shaft. The cam together with the crossbar mechanically trap the shaft inside the stator. This design may be considered as an analogue of a rotaxane, with the stator being the analogue of the ring and the camshaft that of the dumbbell-axle.

To achieve mechanical interlocking, we dock the camshaft first onto one stator unit before closing up the full bearing (Fig. 1F). We achieve this by hybridization of four staple strands protruding from the stator unit to a complementary single-stranded scaffold segment of the camshaft (Fig. 1F, top). We then add the other two stator units, which dock to each other via shape-complementary features carrying sticky ends (Fig. 1F, bottom). Once the complete DNA rotor complex is formed, we release the camshaft from its docking site via toehold-mediated strand displacement. To this end, invader strands are added that remove the initial strand linkages between camshaft and stator unit (Fig. 1G, top). Due to the mechanical interlocking, the camshaft remains sterically trapped in the stator (Fig. 1G, bottom), constrained to a rotary degree of freedom.

### Cryo EM analysis of rotor structure and rotary motions

We self-assembled the components of our rotary mechanism using previously described methods (29) and determined suitable folding conditions using gel electrophoresis (Fig. S4). Trimerization of the stator and also tetramerization of the complete rotary mechanism proceeded robustly (see Methods, Fig. S5). We analyzed the folded structures via negative stain transmission electron microscopy (Fig. S6), and we also determined 3D cryo EM density maps for each of the four DNA origami units of our complex (Fig. 2A-D, Fig. S7-S10), for the trimeric stator lacking the camshaft (Fig. 2E-F, Fig. S11), and for the fully assembled tetrameric rotary mechanism including the camshaft (Fig. 2H, Fig. 3, Fig. S12-S15).

**Figure 2.**
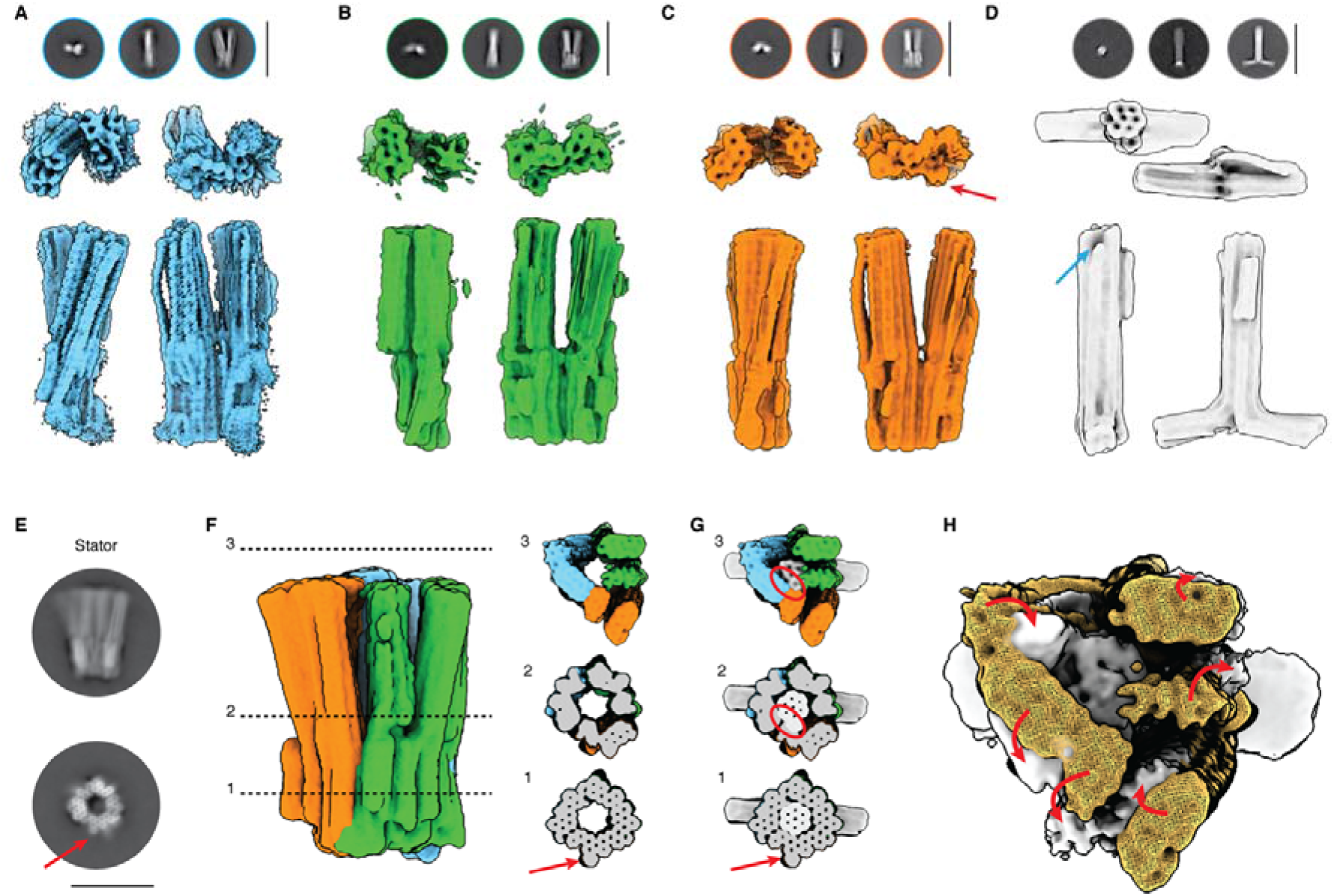
Cryo EM analysis of the components. **(A-D)** Representative 2D class averages (top) and 3D electron density map (bottom), determined from cryo EM micrographs of the three stator units and the camshaft. Red arrow: asymmetric feature on the stator unit 3. Blue arrow: single stranded scaffold segment for successive binding to the stator units. See Fig. S7-S10. Scale bars: 100 nm. **(E, F)** Left to right: 2D class averages, 3D cryo EM map and cross-sectional slices of the 3D EM map of the stator assembled without camshaft. Red arrow indicates the asymmetric feature on stator unit 3. See Fig. S11. Scale bar: 50 nm. **(G)** Slices as in (F), but when the 3D cryo EM map from (D) is placed into the map of the empty stator from (F). Red circles highlight clashes of the camshaft inside the empty stator. **(H)** 3D cryo EM map of the empty stator (yellow) superimposed with the cryo EM map that we determined for one variant of the fully assembled mechanism with the camshaft bound to one of the stator units. Red arrows indicate the deformations in the stator that are necessary to make room for the camshaft.

**Figure 3.**
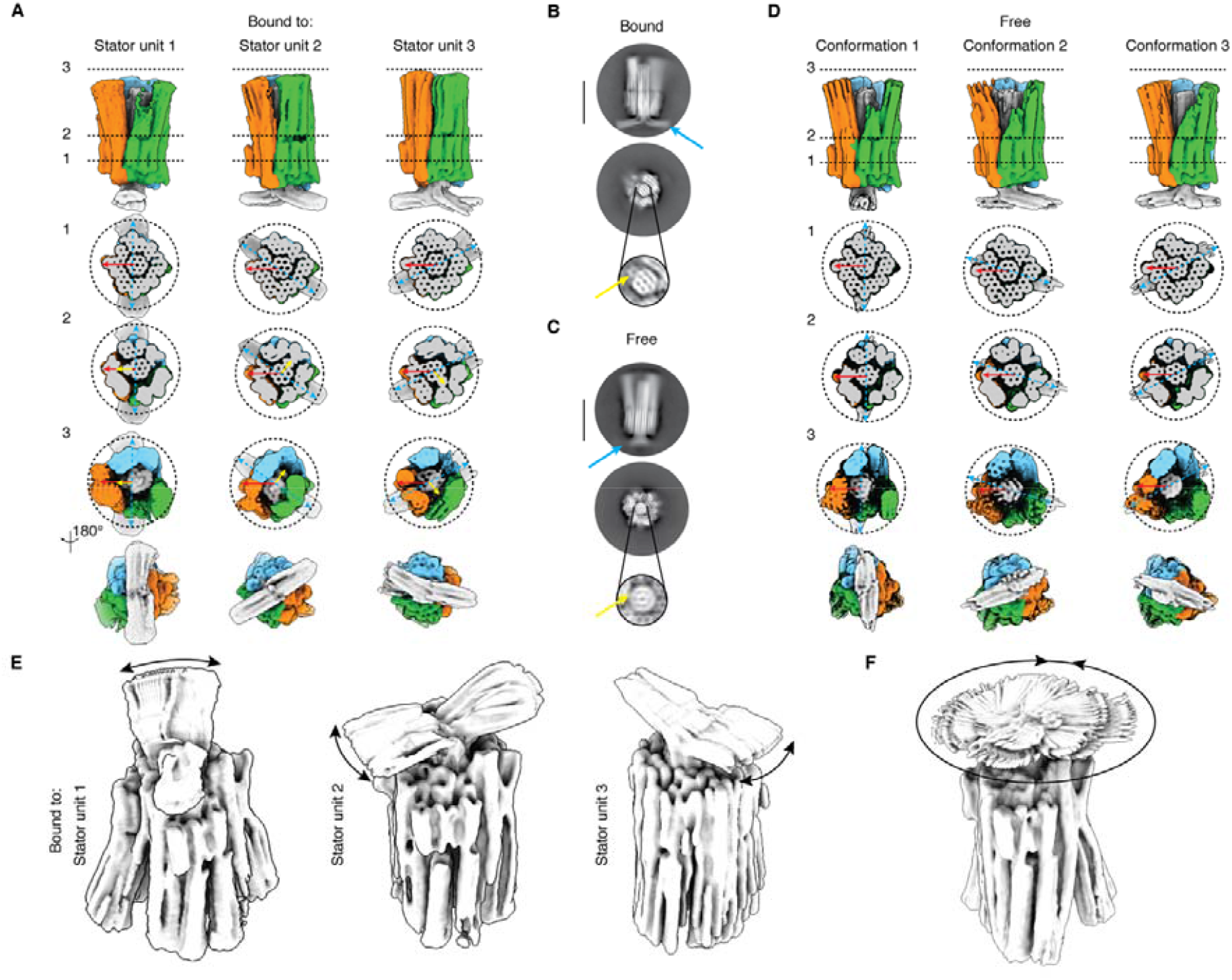
Cryo EM analysis of the whole apparatus. **(A)** Top: side view of the 3D cryo EM maps determined separately for three variants of the mechanism, with the shaft docked to the stator unit 1, 2 or 3, respectively. See Fig. S12-14. Bottom: cross-sectional slices through the maps. Red, blue, yellow arrows highlight the asymmetric feature of the stator, the T-crossbar, and the cam, respectively. **(B)** 2D class averages of the mechanism with the shaft bound to stator unit 1. Blue and yellow arrows highlight crossbar and cam of the shaft, respectively. Scale bar: 50 nm. **(C)** 2D class averages of the mechanism determined from a sample where the shaft was released from the docking site. Blue and yellow arrows point to where crossbar and cam would be expected to be located, but the features are blurred. Scale bar: 50 nm. **(D)** Top: side views of three representative 3D classes discovered in the cryo EM micrograph data determined from the same sample with the released camshaft. Bottom: cross-sectional slices through the 3D classes. Red arrows indicate the asymmetric feature of the stator, blue arrows show the position of the camshaft T-crossbar. The cam could not be resolved in the data. See Fig. S15. **(E)** Overlay of frames from movies from multibody refinement analysis of data acquired separately, with the shaft docked to one stator unit 1, 2 or 3, respectively. Black arrows indicate the observed range of rotary motion of the shaft relative to the stator. **(F)** As in E, but for the sample where the shaft was released from its docking site.

### Structures of components

The cryo EM maps determined for the individual stator units agreed well with the design (Fig. 2A-C). The map determined for stator unit 1 had the highest resolution, with regions where the grooves of the constituent DNA double helices can be discerned. The shape-complementary protrusions and recesses used for docking the stator units into a complete bearing can also be discerned. In stator units 1 and 3 (Fig. 2A, 2C, respectively), the two support DNA double helices to which the pawls are anchored can also be distinguished. The asymmetric feature that marks stator unit 3 can also be discerned (Fig. 2C). In the cryo EM map of the camshaft (Fig. 2D), we can recognize the honeycomb pattern, the protruding “cam” helices on the side of the shaft, and the crossbar. On the other end of the central shaft, we observe a small dent, which we assign to the segment of the scaffold that we left single-stranded in order to allow binding the camshaft to one of the stator units via staple strand hybridization.

### Empty stator vs stator with camshaft inside

The 2D class averages (Fig. 2E) and the 3D cryo map (Fig. 2F) that we determined for the empty stator reveal a structurally well-defined bearing whereas the signal from the pawls is more delocalized and fans away from the long axis of the stator. When slicing through the stator 3D map from bottom to top (Fig. 2F, right), at the level of the empty bearing (slice 1) the honeycomb-shaped cross section matches the design closely (Fig. S16). The asymmetric feature on stator unit 3 is clearly visible. Cutting slightly above the end of the bearing region (slice 2), detail in the cross section is lost and the pawls diverge. Presumably, the loss of detail is due to conformational heterogeneity associated with flexing of the pawls. When looking at the empty stator from the top (slice 3), we see that the pawls are rotated and displaced away from their original position near the bearing to such extent that the central opening is now much smaller than near the bearing further below. When superimposing the map determined separately for the shaft on the map determined for the empty stator (Fig. 2G), we see that the shaft fits well inside the central bore of the stator in the bearing (slice 1), whereas the camshaft and the stator maps show substantial overlap (steric clashes) in the region of the pawls (slices 2 and 3).

In order to accommodate the camshaft into the central bore of the stator, the pawls of the stator therefore need to be pushed outwards. Indeed, in the cryo EM map that we determined from a variant of the entire rotary mechanism with the camshaft docked into the stator (Fig. 2H, Fig. 3), the pawls of the stator are displaced and rotated away from the conformation that they had in the empty stator.

### Camshaft fixed in different orientations

We prepared three distinct variants of the complete rotary mechanism with inserted shaft, in which the camshaft is initially fixed by staple strand linkages to stator units 1, 2, and 3, respectively. These variants thus realize three different, fixed orientations of the camshaft relative to the surrounding stator. These variants allowed us to investigate whether placing the camshaft at different orientations into the stator causes distinct shapes of the stator. We determined 3D cryo EM maps for each of these variants (Fig. 3A). In the resulting maps we could discriminate the asymmetric feature present in the stator unit 3 and used it to assign the stator unit identities and to align the stator orientation. The camshaft indeed assumes three distinct positions inside the stator, rotated by 120°. These orientations can be discerned by comparing the orientation of the asymmetric feature in the stator (Fig. 3A, red arrows) relative to those of the camshaft crossbar (Fig. 3A, blue arrows). We note that the protruding cam on the camshaft can also be discerned in each of the three maps (Fig. 3A, yellow arrows). As designed, the cam is always oriented at 90° relative to the camshaft crossbar orientation. Inspection of cross-sectional slices of the three cryo EM maps with fixed shaft reveals that at the level of the bearing (slice 1), the structures of the variants are all very similar. By contrast, at the level of the pawls, and seen from the top (slices *2, 3*), the maps differ. One can clearly see that the shape of the voids between the six pawls and the central shafts depend on how the shaft is oriented relative to the stator.

### Rotary motion

We prepared the mobile mechanism by releasing the camshaft from the docking site by toehold-mediated strand displacement (see Methods). The conditions for successful release were optimized and validated using electrophoretic mobility analysis performed with incomplete stators (Fig. S17). We acquired cryo EM data of the rotary complex with the camshaft now free to rotate. 2D class averages already reveal crucial differences between those obtained from rotor complexes with a fixed camshaft (Fig. 3B) and those obtained from the sample with presumably mobile camshaft (Fig. 3C). For instance, the honeycomb cross section of the shaft, the cam, and the horizontal crossbar are all clearly visible in the data with fixed shaft (Fig. 3B). On the other hand, these details are blurred in the sample with the released camshaft. The camshaft cross section also now appears as a rotationally averaged version of a honeycomb (Fig. 3C). These images thus suggest that the camshaft is indeed rotating inside of the stator.

We acquired cryo EM micrographs of the rotary mechanism with the mobile camshaft. We analyzed the data set first using 3D classification, which revealed three dominant, structurally distinct 3D classes in the data set (Fig. 3D). We aligned the stator of the maps using the asymmetric feature of stator unit 3 (Fig. 3D, red arrows). Each of the three 3D classes shows a different orientation of the T-crossbar of the shaft (Fig. 3D, blue arrows) relative to the asymmetric feature of the stator. The shaft-to-stator orientations are very similar to those that we prepared by fixing the camshaft to the stator with staple strands (Fig. 3A). However, the cam of the camshaft could not be resolved in the 3D classes. The three 3D classes could thus each contain a mixture of particles featuring camshafts in two different orientations, rotated by 180°.

We also employed multibody refinement to investigate the motion of the camshaft relative to the stator (30). To this end, the components are treated as rigid bodies that can move independently from each other. Using principal component analysis on the relative orientations of the bodies over all particle images in the data set, movies for the important motions in the data can be computed. To illustrate these motions in a still image, we superimposed the frames of the resulting movies (Fig. 3E, F). For the three samples where we fixed the camshaft by strand linkages to the stator, we see that the dominant motion of the camshaft is restricted to some rotary wiggling with a ∼ 20° range (Fig. 3E). By contrast, in the sample with the free camshaft, the rotations of the camshaft cover the entire 360° range (Fig. 3F). Together, the results from 3D classification and multibody analysis indicate that the camshaft can freely rotate inside the stator and that it has at least three, possibly six, preferred orientations.

### Analysis of rotary motions in real-time and with simulations

#### Single-particle fluorescence measurements

We used total internal reflection fluorescence microscopy (TIRFM) to study the dynamical behavior of our rotary complex. To anchor the stator to a glass slide covered by PEG-silane and biotin, we extended stator unit 1 with a helix bundle (6hb) domain at the top of one of the pawls (Fig. 4A, Fig. S18-S19). The 6hb was also labeled with 10 fluorescent dyes (cy5) to detect the position of the stator. Protruding from the 6hb are eight DNA adapter strands to which we hybridized biotinylated DNA strands (Fig. S20) that anchor the stator in a multivalent attachment to the slide via biotin-neutravidin-biotin bridges. The multivalent binding is crucial to suppress rotations of the entire mechanism on the glass slide. We also extended the T-crossbar of the camshaft with a 10-helix bundle lever arm featuring 10 fluorescent dyes (cy3) at its tip, resulting in a ∼ 290 nm long “pointer” (Fig. 4A, Fig. S18, S21) that amplifies and also slows down the motions of the camshaft due to friction with the solvent to facilitate tracking the camshaft motions in real-time.

**Figure 4.**
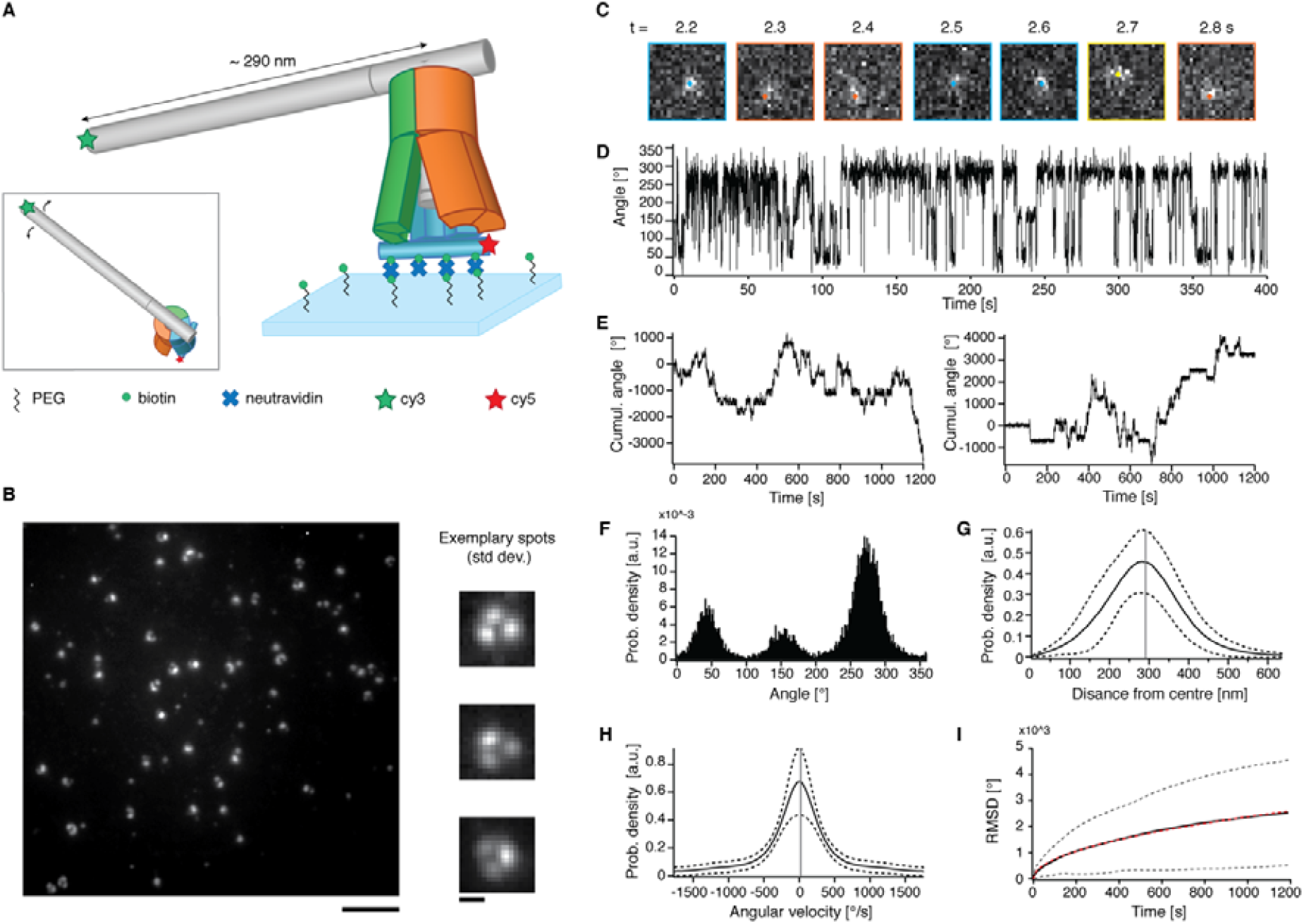
Dynamic analysis of the rotary mechanism via single-particle TIRFM. **(A)** Sketch of the experimental setup: the mechanism is fixed to the cover slide via biotin-neutravidin-biotin bridges. The crossbar of the camshaft is lengthened to approx. 290 nm. Inset: top view. **(B)** Typical field of view image (left, scale bar: 5 µm) and standard deviations of the mean integrated over entire movie for exemplary particles (right, scale bar: 600 nm). **(C)** Exemplary sequence of frames of a typical particle jumping between three preferred positions (blue, orange and yellow). **(D)** Exemplary angle-time trace of a single particle. **(E)** Cumulative angle over time for two typical particles. **(F)** Probability density distribution for lever orientations computed from angle-time trace of a typical particle. **(G)** Solid line: Distribution of the measured distance of lever arm tip from center averaged over N=212 particles. Dashed lines: standard deviation. Vertical gray line: weighted average of the distance from center. **(H)** Solid line: angular velocity distribution computed from N=212 particles. Dashed lines: plus/minus standard deviation. Vertical gray line indicates the weighted average of the angular velocity (which is zero, since there is no directional bias). **(I)** Solid line: average angular root mean square deviation over time from N=212 particles. Dashed lines: plus/minus standard deviation. Red dashed line: fit using 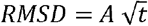.

#### Rotary random walks

Imaging of rotary mechanisms with released camshafts revealed particles performing rotary random walks in addition to stationary particles (Fig. 4B). The rotary particles could potentially reflect situations where the whole complex including the stator is rotating around a faulty connection with the surface. To test for this possibility, we designed a 160 base pair long extension to the 6hb domain protruding from the stator unit 1 (Fig. S22-S23). We thus obtain two pointers that allow tracking simultaneously the motions of the camshaft and of the stator, respectively. With this setup, we confirmed that the rotating particles indeed reflect camshaft motions relative to stator, while the stator remains fixed. In another control experiment with a sample where the camshaft was fixed to the stator by strand linkages, we observed a negligible fraction of particles exhibiting rotary motion. We conclude that the rotary particles seen for the sample with released camshaft indeed reflect motions of the camshaft inside of the stator.

We used super resolution centroid tracking (*31*) (Fig. 4C) to compute single particle angular orientation trajectories (Fig. 4D). These traces typically featured stepwise jumps between three different levels. Since the rotation occurs in thermal equilibrium powered by random Brownian motion, no effective directional bias is expected and is also not observed (Fig. 4E). The particles preferentially populate three main orientations separated by 120° (Fig. 4F), which matches with the designed three-fold symmetry of the stator and with the three preferred orientations of the shaft seen with cryo EM. The mean distance of the moving centroid to the center of movement computed for each particle was ∼286 nm (Fig. 4G), which corresponds well to the expected 290 nm (Fig. 4A). The angular velocity distribution averaged over all measured rotary particles (Fig. 4H) has an approximately Gaussian shape and is centered at 0° as expected for a random Brownian diffusion process without directional bias. Particles can apparently rotate with speeds in the range of up to 3 revolutions per second. The root mean square deviation (RMSD) of the angular displacements grows with the square-root of time, also in accordance with normal diffusion (Fig. 4I).

### Allosteric coordination of rotor and stator motions

We designed and self-assembled five additional variants of the stator in which we altered the flexibility of the pawls (Fig. 5A-F, Fig. S24-S29). All variants were analyzed dynamically with multi-resolution MD simulations using mrDNA (*32*) (Supplementary Movie 1). Experimentally, we released the shaft from its docking site in each sample, acquired single-particle fluorescence microscopy movies, and performed centroid tracking of single rotating particles to compute angular RMSD over time traces (Fig. 5G) as described in Fig. 4I. The stator design modifications had strong influence on the rotary mobility of the camshaft. In variant 1, there are no lateral connections between the pawls, and the pawls can therefore flex independently, which is also seen in the mrDNA simulations (Fig. 5A). In variant 2, we additionally deactivated the base stacking contacts between bearing and pawls, which increases pawl flexibility (Fig. 5B). Experimentally we saw that the rotary mobility of this variant increased approximately by a factor of 2 compared to variant 1 with the stiffer pawls (Fig. 5G). In variant 3, which was already characterized in Fig. 4, we coupled the two pawls within each stator unit laterally with strand crossovers so that they move as one unit (Fig. 5C). This design change removes three of six possible “slots” for the camshaft. Interestingly, variant 3 had very similar rotary mobility compared to variant 1. In variant 4, we coupled the pawls along the lateral interface of neighboring stator units with strand crossovers (Fig. 5D). This design change removes the other three possible slots and had drastic influence on the mobility: it completely inhibited rotary motion, meaning there was a negligible fraction of rotating particles in this sample. The observations so far suggest that the camshaft preferentially populates and switches between the three slots located at the interface between stator units, whereas the slots located in between the two pawls per stator unit are not used. In variant 5, instead of direct strand crossovers as in variant 4, we used 25-thymidine-long strand linkages. These linkages not only restore pawl flexibility, but they also push the pawls a bit apart due to the volume taken up by the poly-T linkages (Fig. 5E). Strikingly, this design change completely restored rotary mobility (Fig. 5G). In fact, this variant showed the highest diffusive mobility of all variants which we attributed to the increased distance between the pawls. Finally, in variant 6, all the pawls were tightly connected to each other by lateral staple strand crossovers (Fig. 5F). Consistent with the previous results, this variant did not rotate at all, presumably because the pawls could not give way to the cam and kept it locked in the conformation in which it was docked initially.

**Figure 5.**
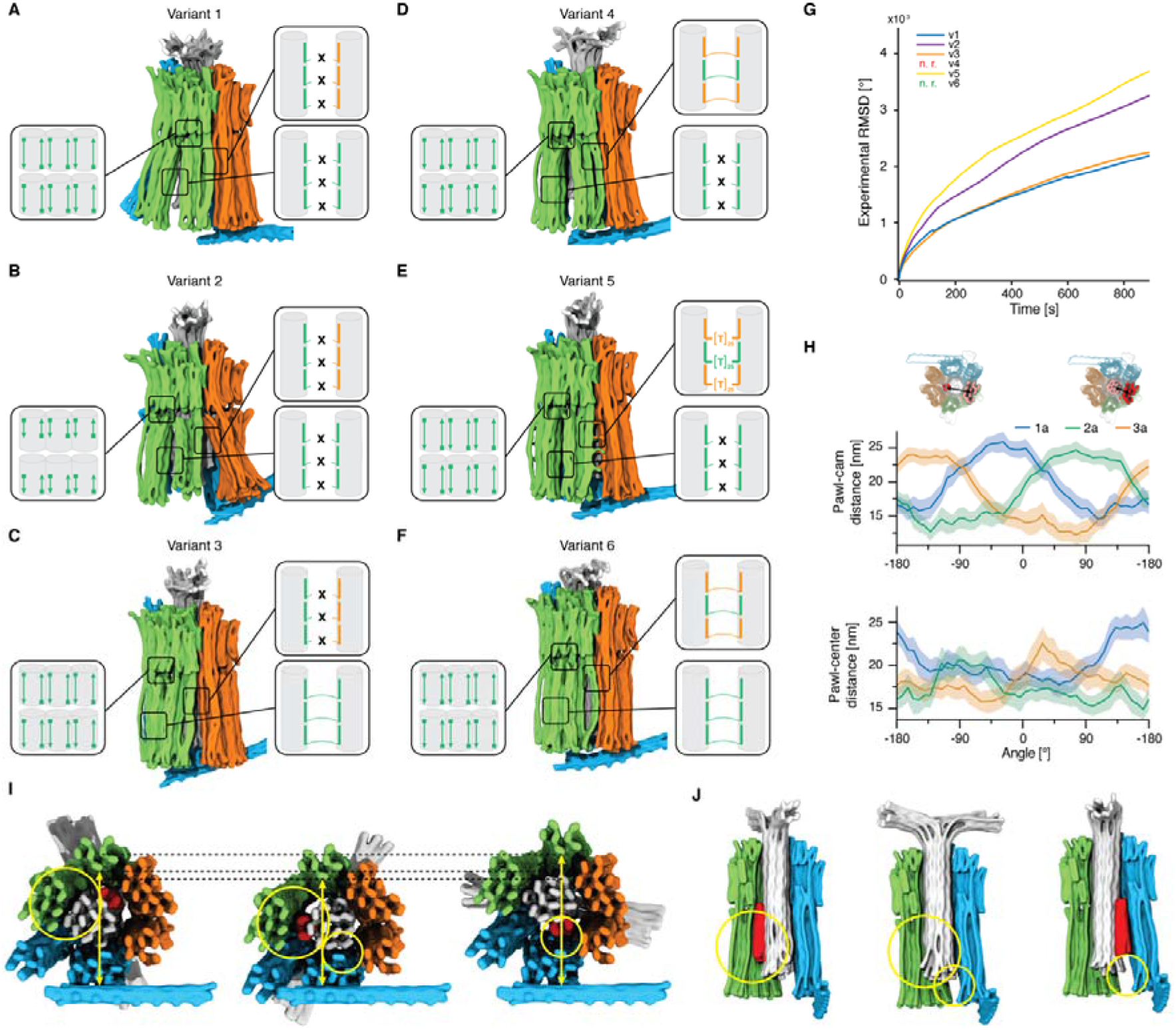
Mechanical coupling between stator and camshaft. **(A-F)** Side views of snapshots from 3D structures predicted for variants of the stator using mrDNA, see Supplementary Movie 1. Insets: schematics of design modifications in stator to influence its mechanical properties. Cylinders represent double stranded DNA helices. **(G)** Experimentally measured average root mean square deviation for variants 1, 2, 3 and 5 from single-particle angle-time traces (see Fig. 4). For variants 4 and 6 no rotation was observed. **(H)** Forced rotation of variant 1. The steered molecular dynamics protocol was applied to a potential acting on the dihedral angle. As the camshaft spins, the cam cyclically approaches each pawl (top), causing it to deform away from the center of the camshaft (bottom). **(I-J)** Average structure from mrDNA simulations of variant 1, extracted from multiple cycles of forced rotation in top (I) and side (J) views. Big circles: deformations on the stator unit 2; small circles: deformations on stator unit 1. In (J) the stator unit 3 is not shown.

MrDNA simulations of variants 1, 3 and 6 at 5 bp/bead resolution investigated the coupling between the camshaft orientation and the mechanical deformation of the stator. In these simulations, the relative orientation of the camshaft and the stator was enforced via a harmonic potential that acted on the dihedral angle formed by the centers of geometry of the four regions (Fig. 5H). By changing the rest angle of the potential at a constant rate, the shaft was driven to rotate in each direction for at least three complete revolutions. We analyzed the resulting trajectories by binning and averaging the microscopic configurations according to the camshaft angle every 10°, revealing deformation of each pawl as the cam approached. For variant 1, the distance between the cam of the shaft and each subunit had a roughly sinusoidal dependence on the camshaft angle with an amplitude of ∼5 nm and phase offset by ∼120° for adjacent subunits, as expected (Fig. 5H, top row). However, when the cam approached a pawl, it caused the pawl to bend away from the center of the camshaft by ∼5 nm as seen by an increase of the distance between each pawl and the center of the shaft (Fig. 5H, bottom row). Similarly, the angle between each pawl and the adjacent pawls was seen to be maximal as the cam approached the outer pawls, and minimal as it approached the central pawl (Fig. S30). These deformations can also be seen from exemplary simulation snapshots (Fig. 5I-J): when the cam approaches one of the pawls, they are pushed further from the camshaft center, while they relax back into position if the cam is pointing away.

We generalized the simulation analysis to variants 3 and 6, revealing that all variants exhibit qualitatively similar deformations of the pawls despite the different inter-pawl connections (Fig. S30). However, the deformation is diminished by increased coupling of the pawls as implemented in variant 3 and especially variant 6, which did not show any actual rotation in our experiments. Furthermore, the minimum pawl–shaft distance during the rotation cycle of variant 1 is similar to the maximum distance for variant 6, reflecting that the latter does not readily accommodate the cam. In summary, the simulations show that the rotary motion of the cam is tightly coupled to, and coordinated by, the reciprocal deformation of the pawls, as designed. The coordination and reciprocal motion may be appreciated in movies of the forced rotation simulation results for variants 1, 3 and 6, respectively (Supplementary Movies 2 and 3).

## Discussion

In this work, we presented a compliant nanoscale rotary mechanism with a central camshaft surrounded by a stator with programmable stiffness. We used single-particle cryo-electron microscopy to structurally characterize the components, and also the entire mechanism in different states. We also studied the dynamical behavior of the rotary apparatus via total internal reflection fluorescence microscopy and molecular dynamic simulations. The results from structural analysis by cryo EM, the single particle fluorescence imaging, and the simulations all support the following picture: the camshaft can freely rotate inside the stator, but there exist three preferred shaft orientations. These orientations correspond to states with the cam snapped into slots located at the boundaries between the stator units.

The three preferred states for the cam are defined mechanically, meaning that the camshaft is pressed into the slots by the forces exerted by the surrounding stator. This is a crucial difference to previously reported nanomachines, where conformational states were defined via direct chemical bonds. The mechanical snapping into place now enables regulation at a distance. In our mechanism, the pawls of the surrounding stator must deform to allow rotation of the camshaft and escape from the mechanical slots, as visualized directly by the simulated trajectories. Such deformations occur in our mechanism in a thermally activated fashion, giving rise to Brownian rotary diffusion. Through targeted design changes, we made some versions of the stator less flexible. As a consequence, the rotary movements of the shaft became slower, even stalling the camshaft in two design variants. Together, these experiments demonstrate an allosteric coupling between the orientation of the camshaft and the mechanical deformation of the surrounding bearing and between the rotary motion of the shaft and the reciprocal open/close transitions in the stator.

Our mechanism could provide a framework for creating artificial nanomachines that, similarly to their counterparts in the macroscopic world, can operate through coordinated motion of their components. For example, it could be of interest to direct the opening/closing transitions in the stator by the consumption of chemical fuel (*6-12*) to create a chemically fueled rotary nanomotor. Likewise, due to microscopic reversibility, it is conceivable that such a system could potentially be reversed and used for uphill chemical synthesis, as in ATP synthase, by applying mechanical torque to the central shaft, thus creating a chemical generator. In that pursuit, the coordinated motions of the pawl and the camshaft, or new variants of it, could be employed to cyclically bring reactants into close proximity. All of these applications require creation of intricately shaped components and their assembly into a functional mechanism. Our work shows a route for how such tasks can be accomplished but also highlights the challenges involved in imparting the desired functionality on such ultraminiaturized molecular mechanisms. We expect that the realization of more complex artificial machinery will go hand in hand with further improvements in analyzing continuous molecular motions by cryo EM (*30, 33*) and with improved predictive computational approaches (*32, 34*).

## Acknowledgements

This work was supported by a European Research Council Consolidator Grant to H.D. (GA no. 724261), the Deutsche Forschungsgemeinschaft through grants provided within the Gottfried-Wilhelm-Leibniz Program and the SFB863 TPA9 Project ID 111166240 (to H.D.). A.A. and C.M.M. acknowledge support through National Science Foundation (USA) under grant DMR-1827346 and the National Institutes of Health under grant P41-GM104601. Supercomputer time was provided through Leadership Resource Allocation MCB20012 on Frontera.

We thank Massimo Kube and Fabian Kohler for helpful discussions on the cryo EM reconstructions, Anna-Katharina Pumm and Wouter Engelen for support with the MATLAB script and the fluorescence microscopy experiments, and Alexander Koch for auxiliary experiments.

## Author contributions

H.D. designed the research. E.B. and T.D. performed the research. M.N.H. provided the custom scaffold. C.M.M. and A.A. performed the simulations. E.B., C.M.M., H.D. and A.A. prepared the figures and wrote the manuscript. All authors have given approval of the final version of the manuscript. The authors declare no conflict of interest.

