## Supplementary Information for "A nanoscale reciprocating rotary mechanism with coordinated mobility control"

**Content**

Materials and Methods

Supplementary Figures S1-S30

### Materials and Methods

##### Design of the DNA origami nanostructures

All structures were designed using cadnano0.2 (*1*). The central shaft was folded from a 7560-bases long scaffold, while the stator units and the lever arm were folded from an 8064-bases long scaffold. The modified stator units 1 for TIRFM measurements were folded from a 9072-bases long scaffold. The 6 helix-bundle bound to the stator unit 1 was folded from a 2873-bases long scaffold (*2*).

##### Folding of the DNA origami nanostructures

The folding reaction mixtures contained 40 nM scaffold and 160 nM (for structures with p9072 scaffold) or 200 nM (for the other structures) staples (Eurofins MWG, IDT). The folding buffer included 5 mM TRIS, 1 mM EDTA, 5 mM NaCl and 10 to 20 mM $MgCl_2$. The folding solutions were thermally annealed using TETRAD (MJ Research, now Biorad) thermal cycling devices. The reactions were left at 65°C for 15 minutes and were subsequently subjected to a thermal ramp from 60°C to 44°C (1°C/hour). The folded structures were stored at room temperature.

##### Purification and concentration of the DNA origami nanostructures

After folding, all samples were purified via PEG purification, ultrafiltration or physical extraction from agarose gel. For concentrating the monomers, ultrafiltration was used, while ultracentrifugation was used for concentrating the polymers. All procedures were performed as previously described (*3*).

##### Assembly of the complex

For the dimerization between stator unit 1 and shaft, a 1:1 solution of the monomers was mixed with 20 mM $MgCl_2$ and left at 40°C for 16-24 hours. The other stator units and the lever arm were added in stoichiometric conditions at 40 mM $MgCl_2$ and left at 40°C for 1 day/monomer addition. To set the central shaft free to rotate, invader strands were added in excess (2x for cryo EM analysis, 4x for TIRFM analysis) and the reaction mixture was left in a shaker for 16-24 hours at room temperature.

##### Agarose gel analysis of the DNA origami nanostructures

The DNA nanostructures were electrophoresed on 2% agarose gels containing 0.5x TBE and $MgCl_2$ at different concentrations: typically, for monomers and dimers 5.5 mM $MgCl_2$ was used, while 22 to 33 mM $MgCl_2$ was necessary for higher order structures. For the latter samples, the temperature was kept at around 10°C with a cooling system. The gels were scanned using a Typhoon FLA 9500 laser scanner (GE Healthcare) at a resolution of 50 µm/pixel.

##### Negative stain transmission electron microscopy

Purified structures were adsorbed onto glow-discharge Cu grids with carbon support (in house production and Science Services, Munich) and stained with a 2% aqueous uranyl formate solution containing 25 mM NaOH. Samples were incubated for different time lengths depending on the concentration. In general, structures with concentrations in the order of tens of nM were incubated for 30 s, while lower concentrated samples (5 nM or below) were incubated for 5 to 10 minutes.

Images were acquired using a Philips CM100 operating at 100 kV or a Tecnai 120 (Thermo Fisher Scientific) at 120 kV. Negative stain 2D class averages were computed using RELION (*4*) without CTF correction.

**Cryo electron microscopy sample preparation**

For cryo EM analysis of the monomers, the samples were folded and ultrafiltrated as previously described (*3*) until reaching concentrations around 1 μM.

For higher order structures, monomers were folded and gel extracted in order to ensure to have only the right monomeric species. The samples were ultrafiltrated for increasing the concentration. The monomers were polymerized as described in the previous paragraph. The polymers were ultracentrifuged at 45k RPM for 30 min at 25°. The supernatant was pipetted away leaving only a small volume of sample with the desired high concentration. If needed, the invader strands were added at a 2x excess to binding site for 16-24 hours at room temperature.

The samples were incubated onto glow-discharged C-flat 1.2/1.3, C-flat 2/1 or lacey carbon with ultrathin carbon support grids and plunged-frozen using a Vitrobot Mark IV (Thermo Fisher Scientific). The parameters used are given in the table below (SU = stator unit; CS = camshaft; B SU = camshaft bound to stator unit; Free = camshaft set free to rotate; LCS = lacey carbon with ultrathin carbon support).

| **Sample** | SU1 | SU2 | SU3 | CS | stator | B SU1 | B SU2 | B SU3 | Free |
|---|---|---|---|---|---|---|---|---|---|
| **Volume [μL]** | 4 | 4 | 4 | 3 | 4 | 4 | 3-4 | 3-4 | 4 |
| **Conc. [nM]** | 1800 | 1400 | 780 | 129 | 80 | 3-100 | 40-85 | 65-92 | 35-93 |
| **Humidity [%]** | 100 | 100 | 100 | 100 | 100 | 100 | 100 | 100 | 100 |
| **Temp. [°C]** | 20 | 20 | 20 | 20 | 20 | 20 | 20 | 20 | 20 |
| **Wait t [s]** | 0 | 0 | 0 | 60 | 0-5 | 0-600 | 0 | 0 | 0-15 |
| **Blot t [s]** | 1 | 2 | 2 | 0 | 1-2 | 0-3 | 0-2 | 0-2 | 0-3 |
| **Blot force [mm]** | 0 | 0 | 0 | 1 | 1-2 | 0-1 | 0-2 | 1-2 | 1-2 |
| **Drain t [s]** | 0 | 0 | 0 | 3 | 0 | 0 | 0 | 0 | 0 |
| **Grid type** | C-flat 1.2/1.3 | C-flat 1.2/1.3 | C-flat 1.2/1.3 | C-flat 1.2/1.3 | C-flat 1.2/1.3 | C-flat 1.2/1.3, LCS | C-flat 1.2/1.3 | C-flat 1.2/1.3 | C-flat 1.2/1.3 |
| **Number of grids** | 1 | 1 | 1 | 1 | 2 | 9 | 6 | 5 | 8 |

##### Cryo electron microscopy image acquisition

The camshaft was manually imaged using a Tecnai 120 (Thermo Fisher Scientific). All the other samples were imaged automatically with a Titan Krios (Thermo Fisher Scientific). Imaging parameters are listed in the table below.

| Sample | SU1 | SU2 | SU3 | CS | stator | B SU1 | B SU2 | B SU3 | Free |
|---|---|---|---|---|---|---|---|---|---|
| **Microscope** | Titan | Titan | Titan | Tecnai 120 | Titan | Titan | Titan | Titan | Titan |
| **Voltage [kV]** | 300 | 300 | 300 | 120 | 300 | 300 | 300 | 300 | 300 |
| **Magnification** | 29kx | 29kx | 29kx | 30kx | 29kx | 37kx | 29kx | 29kx | 29kx |
| **Spot size** | 4 | 4 | 4 | / | 4 | 5 | 4 | 4 | 4 |
| **Defocus [µm]** | -2 | -2 | -2 | -2 | -2 | -2 | -2 | -2 | -2 |
| **Dose [$e^-/A^2$]** | ~40 | ~40 | ~40 | / | ~40 | ~45 | ~50 | ~40 | ~50 |
| **Exposure time [s]** | 2.6 | 2.6 | 2.6 | 1 | 2.6 | 1.5 | 2.6 | 2.6 | 2.6 |
| **Pixel size [A]** | 2.32 | 2.32 | 2.32 | 3.37 | 2.32 | 1.82 | 2.32 | 2.32 | 2.32 |
| **Frames** | 105 | 105 | 105 | / | 105 | 63 | 105 | 105 | 105 |
| **Fraction** | 7 | 7 | 7 | / | 7 | 9 | 7 | 7 | 10 |

##### Cryo electron microscopy image processing

The movies acquired with the Titan were subjected to motion correction using MotionCor2 (*5*). All the micrographs were CTF corrected with CTFFIND4 (*6*). The camshaft particles were picked manually, while for the other samples particles were picked with RELION (*4*) or with crYOLO (*7*). Other processing steps were performed with RELION (RELION 2 for the central camshaft, RELION 3.0 for the other structures). Multiple runs of 2D and 3D classifications were typically performed to exclude incompletely folded particles or particles lying on the carbon film. The 3D classes presenting most features were further refined and post-processed.

For the complex set free to rotate, a 3D classification without alignment was performed after refinement, using multiple maps as references, i.e., the three reconstructions of the structure with the camshaft fixed to the stator units. With this method, three different positions of the central camshaft could be found.

The stator units were further subjected to a MultiBody refinement (*8*), where each of the pawls and the bearing were treated as a separate body. Similarly, the higher order structures were subjected to Multibody refinements where each of the monomers composing the complex was considered a separate rigid body.

##### Assembly for total internal reflection fluorescence microscopy

Monomers were folded and purified using physical gel extraction. The samples were ultrafiltrated for increasing the concentration. The monomers were polymerized as described before. If needed, the invader was added at a 4x excess to binding site for 16-24 hours at room temperature. Biotinylated oligos were incubated with a 32x excess neutravidin and then added to the polymers in an 8x excess to binding site for 1-2h at room temperature. The resulting reaction mixture was gel purified for extracting only the correct species. Sample concentrations varied between 100 and 700 pM. Samples were left at RT for at most 2 days and then imaged at the microscope.

##### Total internal reflection fluorescence microscopy movie acquisition

Microscope cover slides were functionalized with bioPEG as previously described (*9*).
An acrylic glass template containing 4 chambers was sealed to the cover slide with vacuum grease. The chambers were washed with 2xFoB20 buffer (10 mM TRIS, 2 mM EDTA, 10 mM NaCl, 40 mM $MgCl_2$) for 3-4 times. 50 µL samples was incubated for 5 to 20 min depending on the sample concentration. The chamber was washed again 3-5 times with a buffer containing 5 mM TRIS, 1 mM EDTA and 500 to 1000 mM NaCl. Afterwards, the chamber was washed 3 times with imaging buffer containing an oxygen scavenging system (50 mM TRIS pH 8, 1 mM EDTA, 500 mM NaCl, 2 mM Trolox, 0.8% D-glucose, 442 U/ml glucose oxidase, 2170 U/ml catalase) prior to data acquisition. Enzymes, Trolox and glucose were purchased from Sigma Aldrich.
Movies were acquired at room temperature with a custom-built objective-type TIRFM (*10*).
Movies were acquired for 2 to 20 min at a frame rate of 20 frames/s and a laser on time of 5 ms.

##### Total internal reflection fluorescence microscopy movie processing

Movies were drift corrected using a FIJI plugin (NanoJ-Core (*11*)). All successive steps were performed using a custom MATLAB script. Moving particles were manually picked and their frame-by-frame standard deviation was computed. Defective particles or particles showing no motion were sorted out by inspecting the standard deviation images. The remaining particles were further processed by tracking the position of the lever arm in each frame using a virtual window center of mass approach (VWCM). The obtained spots were clustered in 3 groups indicating the three preferred positions of the central camshaft. From the tracked spots, angular positions, RMSD and angular velocities (Ω) were calculated according to:

$$RMSD = \sqrt{\frac{\sum_{i=1}^{N}(\theta_i - \theta_0)^2}{N}}$$

$$\Omega_i = \frac{\theta_i - \theta_{i-1}}{\Delta t}$$

where $N$ is the number of frames, $i$ indicates the *i-th* frame, $\theta_0$ is the angle at t = 0 and Δt indicates the time difference between 2 consecutive frames. A histogram of the angular velocities was calculated.

##### Multi-resolution simulations

Each variant was simulated using the mrDNA multiresolution modeling framework (*12*), first at a resolution of ~5 bp/bead with a 200 fs timestep for 20 µs, followed by simulations at 2-bp/bead resolution and a 40-fs timestep that lasted 80 ns. The higher resolution simulations introduced a local representation of the orientation of the major groove that facilitated construction of an atomic model of each variant. The temperature was held at 295 K. The computational model of DNA–DNA interactions was previously parameterized (*12*) to match the experimentally-measured osmotic pressure in a DNA condensate at 20 mM $MgCl_2$ electrolyte. The stacking interactions between the DNA helices in the bearing region and the pawls were modeled within the mrDNA framework using a custom script that effectively made the two helices contributing to the stacking site a single continuous dsDNA helix, with the azimuthal angle of the helices on both sides of each stacking site being in phase.

Selected variants (1, 3, and 6) were additionally simulated with an applied bias to drive the rotation. The simulations were performed as described above, except a harmonic potential ($k_{spring}$ = 0.5 kcal $mol^{-1}$ $degree^{-2}$) was placed on a dihedral angle formed by the centers of geometry of the following four groups of particles: (i) beads constituting the central 60 bp of each of the six-helices that formed the “cam” of the rotor; (ii) beads in the same 60-bp-thick plane of the cadnano design that formed the 24-helix shaft of the rotor; (iii) a 60-bp-thick slab of beads in the shaft of the rotor near the bearing region separated by 85 bp (center-to-center) from the other group of beads on the shaft; and (iv) a 40-bp thick slab of beads in the center of the second subunit of the stator (between 50 and 90 bp from the bottom edge of the bearing). For each variant, two simulations were performed with the rest angle of the harmonic potential increasing in one and decreasing in the other. The simulations were performed for a sufficiently long period of time to observe three complete rotations in each direction.

### Supplementary Figures

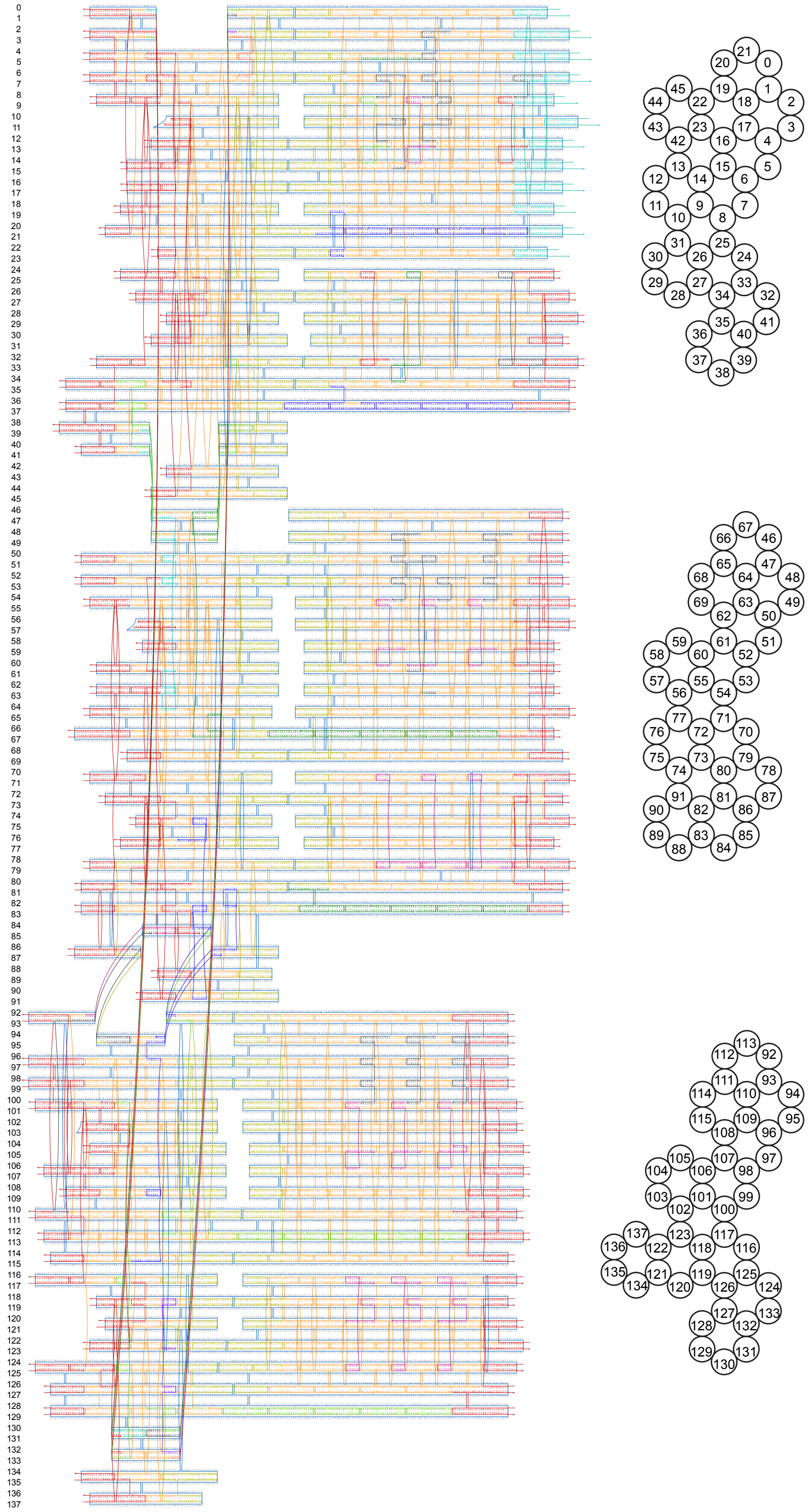

**Figure S1** CaDNAno (*1*) design diagrams (left) and bottom view cross sections (right) of the stator units 1 (top), 2 (middle) and 3 (bottom).

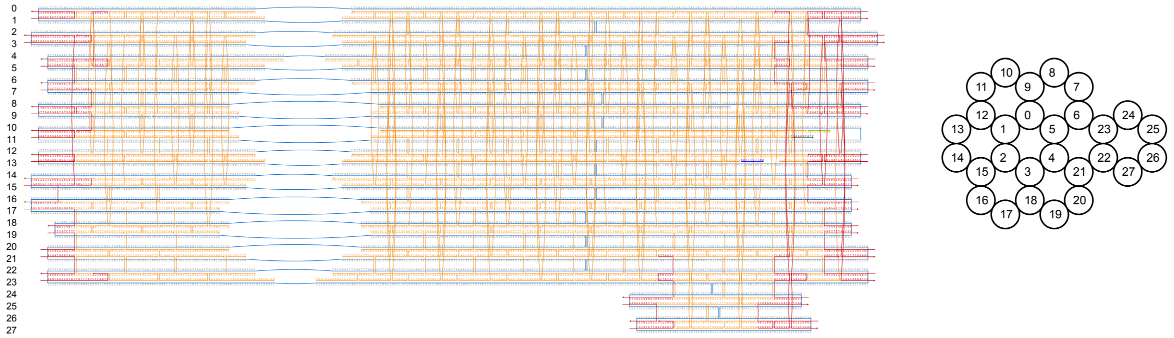

**Figure S2** CaDNAno (*1*) design diagram (left) and bottom-view cross section (right) of the central camshaft.

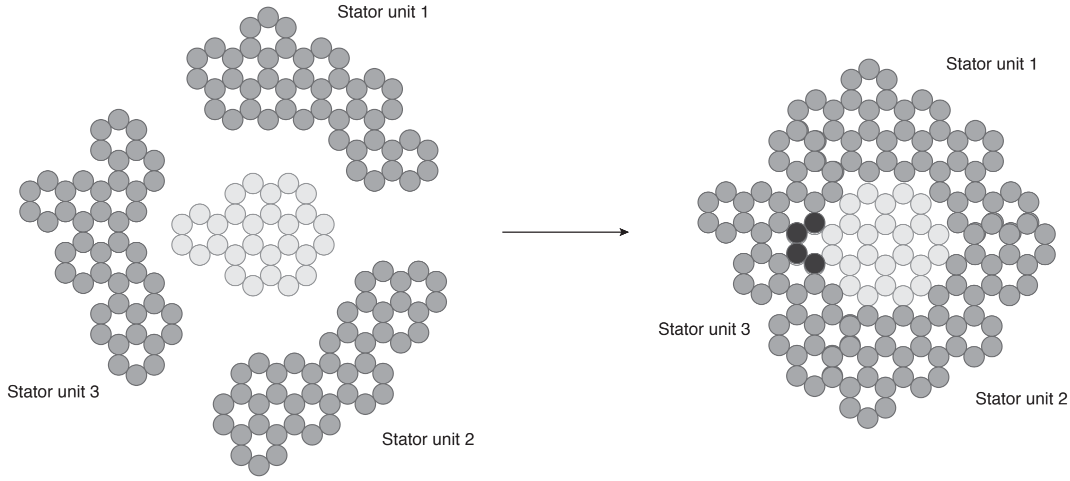


**Figure S3 Schematic representation of the helical cross section of the rotary mechanism.** Each circle represents a DNA double helix. Top view cross sections of the monomers composing the mechanism (left) and cross section of the fully formed mechanism (right). In black the helices of the camshaft that overlap with the helices of the surrounding stator.

**A**

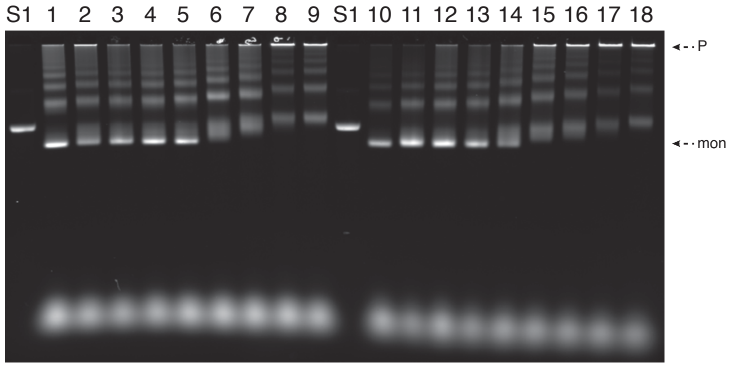


**B**

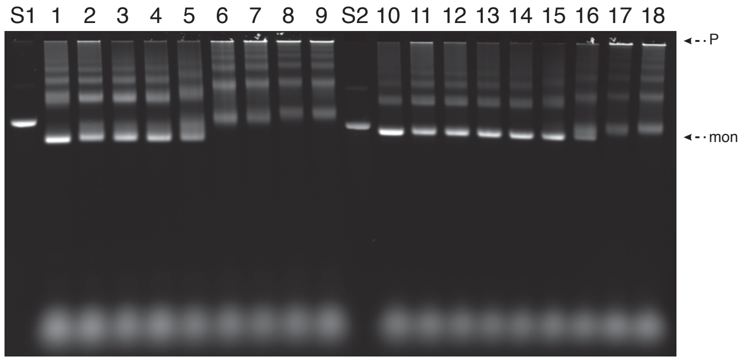


**Figure S4 Temperature folding screens of the monomers. (A)** Laser-scanned photograph of a 2% agarose gel with temperature folding screens of stator unit 1 (1-9) and stator unit 2 (10-18). **(B)** Laser-scanned photograph of a 2% agarose gel with temperature folding screens of stator unit 3 (1-9) and camshaft (10-18). S1: p8064 scaffold; S2: p7560 scaffold. Lanes 1, 10: 60°C-44°C; lanes 2, 11: 50°C-47°C; lanes 3, 12: 52°C-49°C; lanes 4, 13: 54°C-51°C; lanes 5, 14: 56°C-53°C; lanes 6, 15: 58°C-55°C; lanes 7, 16: 60°C-57°C; lanes 8, 17: 62°C-59°C; lanes 9, 18: 64°C-61°C. P: pockets; mon: monomers.

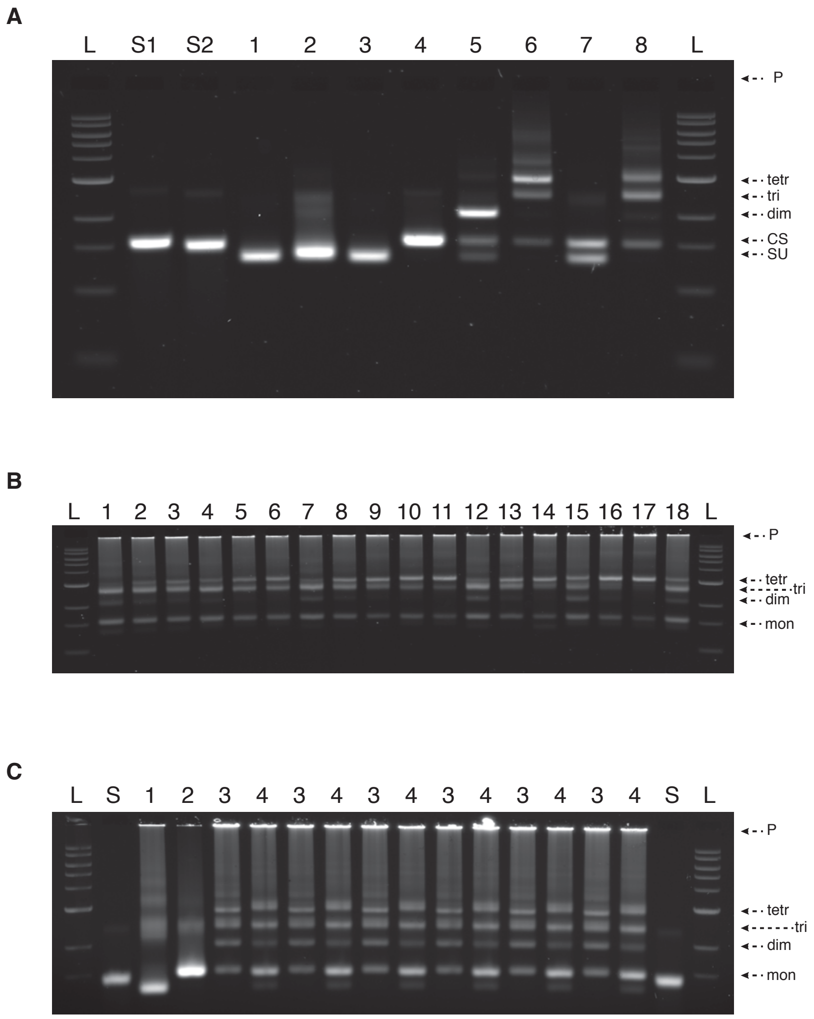


**Figure S5 Assembly of the complex. (A)** Assembling the complex. Laser-scanned photograph of a 2% agarose gel on which the following samples were electrophoresed: L: ladder; S1: p8064 scaffold; S2: p7560 scaffold; lane 1: stator unit 1; lane 2: stator unit 2; lane 3: stator unit 3; lane 4: camshaft; lane 5: stator unit 1 and camshaft dimers; lane 6: tetrameric complex; lane 7: stator unit 1 and camshaft dimers with invader strands added; lane 8: tetrameric complex with invader strands added. **(B)** Assembly screen for the complex with the camshaft bound to different stator units. Laser-scanned photograph of a 2% agarose gel on which the following samples were electrophoresed: L: ladder; lane 1: camshaft bound to stator unit 2, 20 mM $MgCl_2$, 30°C; lane 2: camshaft bound to stator unit 2, 30 mM $MgCl_2$, 30°C; lane 3: camshaft bound to stator unit 2, 40 mM $MgCl_2$, 30°C; lane 4: camshaft bound to stator unit 3, 20 mM $MgCl_2$, 30°C; lane 5: camshaft bound to stator unit 3, 30 mM $MgCl_2$, 30°C; lane 6: camshaft bound to stator unit 3, 40 mM $MgCl_2$, 30°C; lane 7: camshaft bound to stator unit 2, 20 mM $MgCl_2$, 40°C; lane 8: camshaft bound to stator unit 2, 30 mM $MgCl_2$, 40°C; lane 9: camshaft bound to stator unit 2, 40 mM $MgCl_2$, 40°C; lane 10: camshaft bound to stator unit 3, 30 mM $MgCl_2$, 40°C; lane 11: camshaft bound to stator unit 3, 40 mM $MgCl_2$, 40°C; lane 12: camshaft bound to stator unit 2, 20 mM $MgCl_2$, 50°C; lane 13: camshaft bound to stator unit 2, 30 mM $MgCl_2$, 50°C; lane 14: camshaft bound to stator unit 2, 40 mM $MgCl_2$, 50°C; lane 15: camshaft bound to stator unit 3, 20 mM $MgCl_2$, 50°C; lane 16: camshaft bound to stator unit 3, 30 mM $MgCl_2$, 50°C; lane 17: camshaft bound to stator unit 3, 40 mM $MgCl_2$, 50°C; lane 18: camshaft bound to stator unit 3, 20 mM $MgCl_2$, 40°C. **(C)** Difference between the complex when the central camshaft is bound and not bound. Laser-scanned photograph of a 2% agarose gel on which the following samples were electrophoresed: L: ladder; S: p7560 scaffold; lane 1: stator unit 3; lane 2: camshaft; lanes 3: complex with the camshaft bound to stator unit 1; lanes 4: complex with camshaft free to rotate. P: pockets; tetr: tetramers; tri: trimers; dim: dimers; mon: monomers; CS: camshaft; SU: stator units.

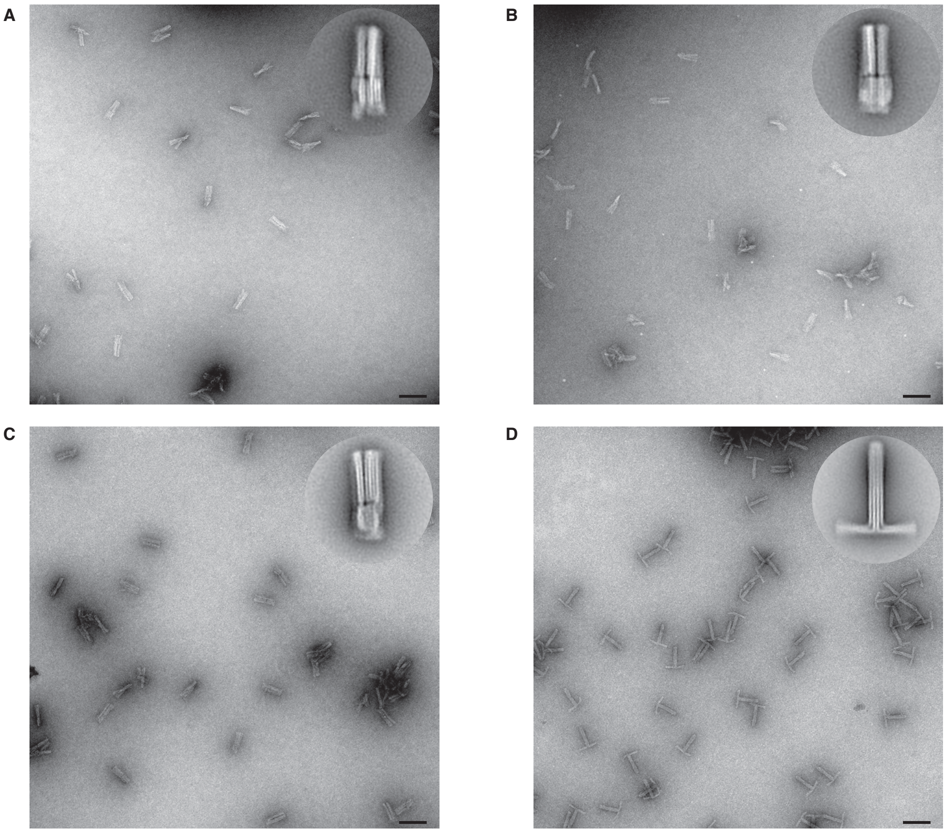

**Figure S6 Negative stain electron micrographs of the components of the rotary mechanism. (A)** Stator unit 1. **(B)** Stator unit 2. **(C)** Stator unit 3. **(D)** Camshaft. Scale bars 100 nm. Insets: 2D class averages of the monomers.

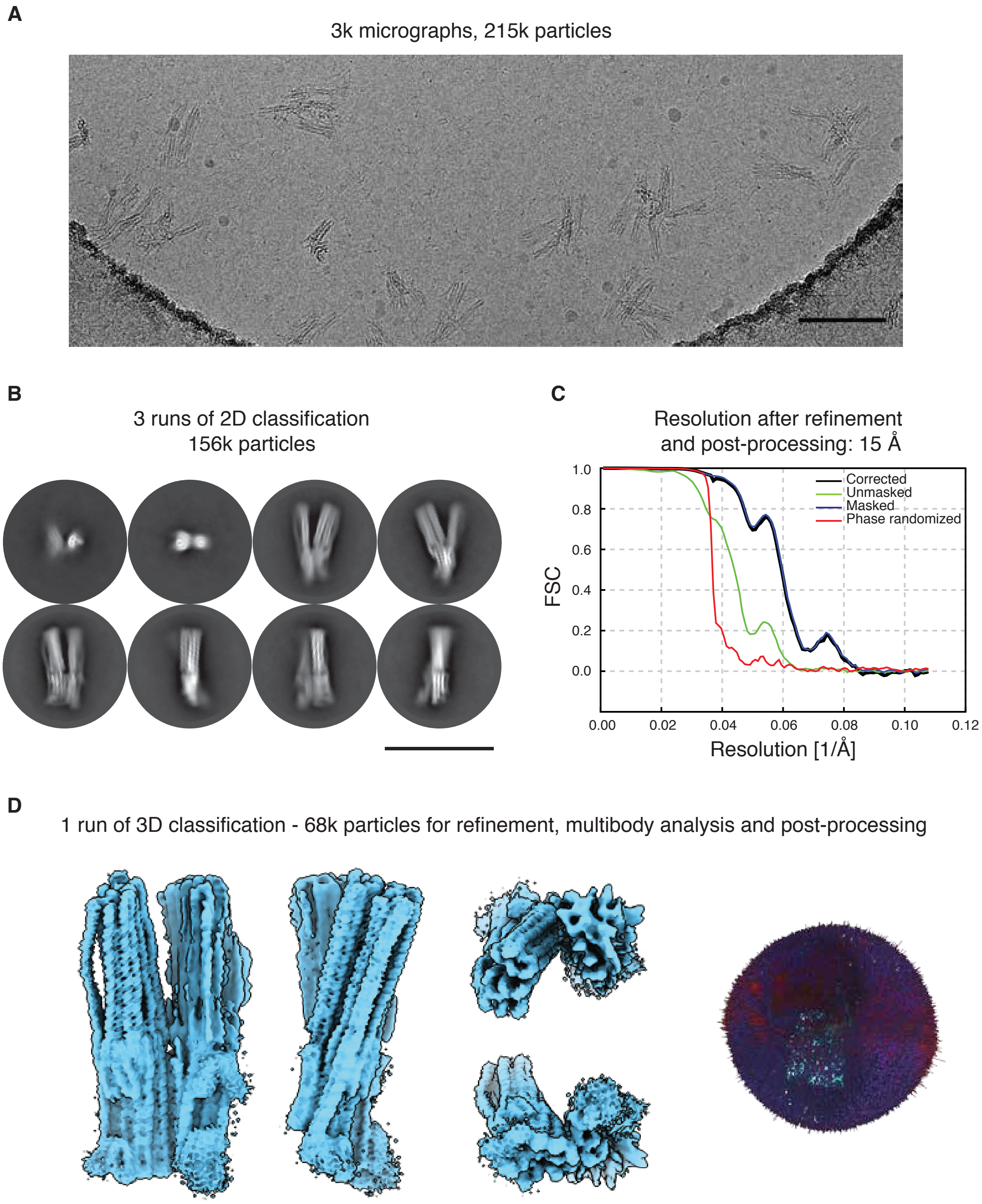


**Figure S7 Cryo EM reconstruction of stator unit 1. (A)** Exemplary motion-corrected and dose-weighted micrograph. Scale bar 100 nm. **(B)** Exemplary 2D class averages showing the particles in different views. Scale bar 100 nm. **(C)** Fourier Shell Correlation (FSC) of the refined map. **(D)** Composite map from a MultiBody job in different orientations (left) and three-dimensional histogram of the particle orientations (right).

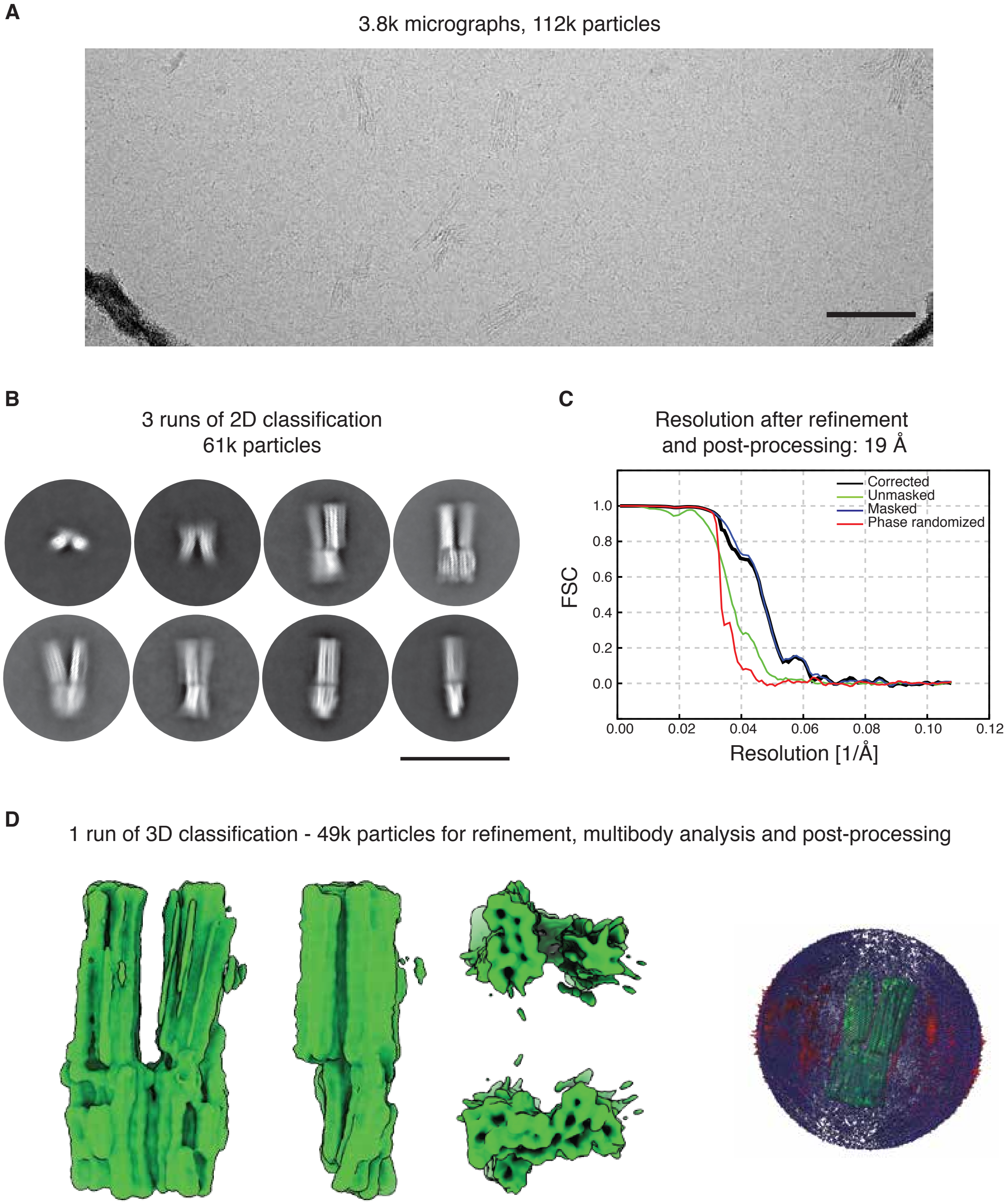


**Figure S8 Cryo EM reconstruction of stator unit 2. (A)** Exemplary motion-corrected and dose-weighted micrograph. Scale bar 100 nm. **(B)** Exemplary 2D class averages showing the particles in different views. Scale bar 100 nm. **(C)** Fourier Shell Correlation (FSC) of the refined map. **(D)** Composite map from a MultiBody job in different orientations (left) and three-dimensional histogram of the particle orientations (right).

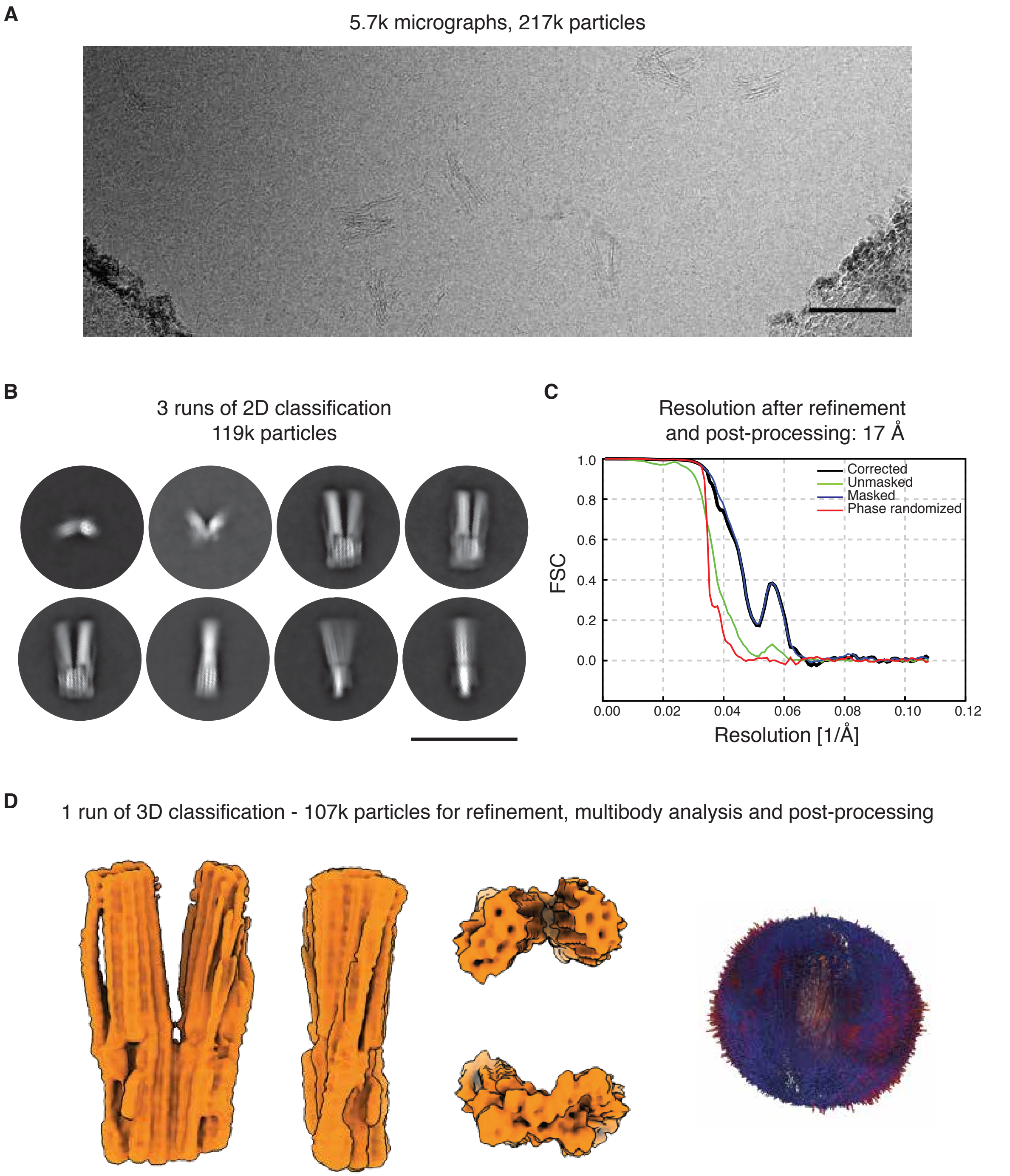


**Figure S9 Cryo EM reconstruction of stator unit 3. (A)** Exemplary motion-corrected and dose-weighted micrograph. Scale bar 100 nm. **(B)** Exemplary 2D class averages showing the particles in different views. Scale bar 100 nm. **(C)** Fourier Shell Correlation (FSC) of the refined map. **(D)** Composite map from a MultiBody job in different orientations (left) and three-dimensional histogram of the particle orientations (right).

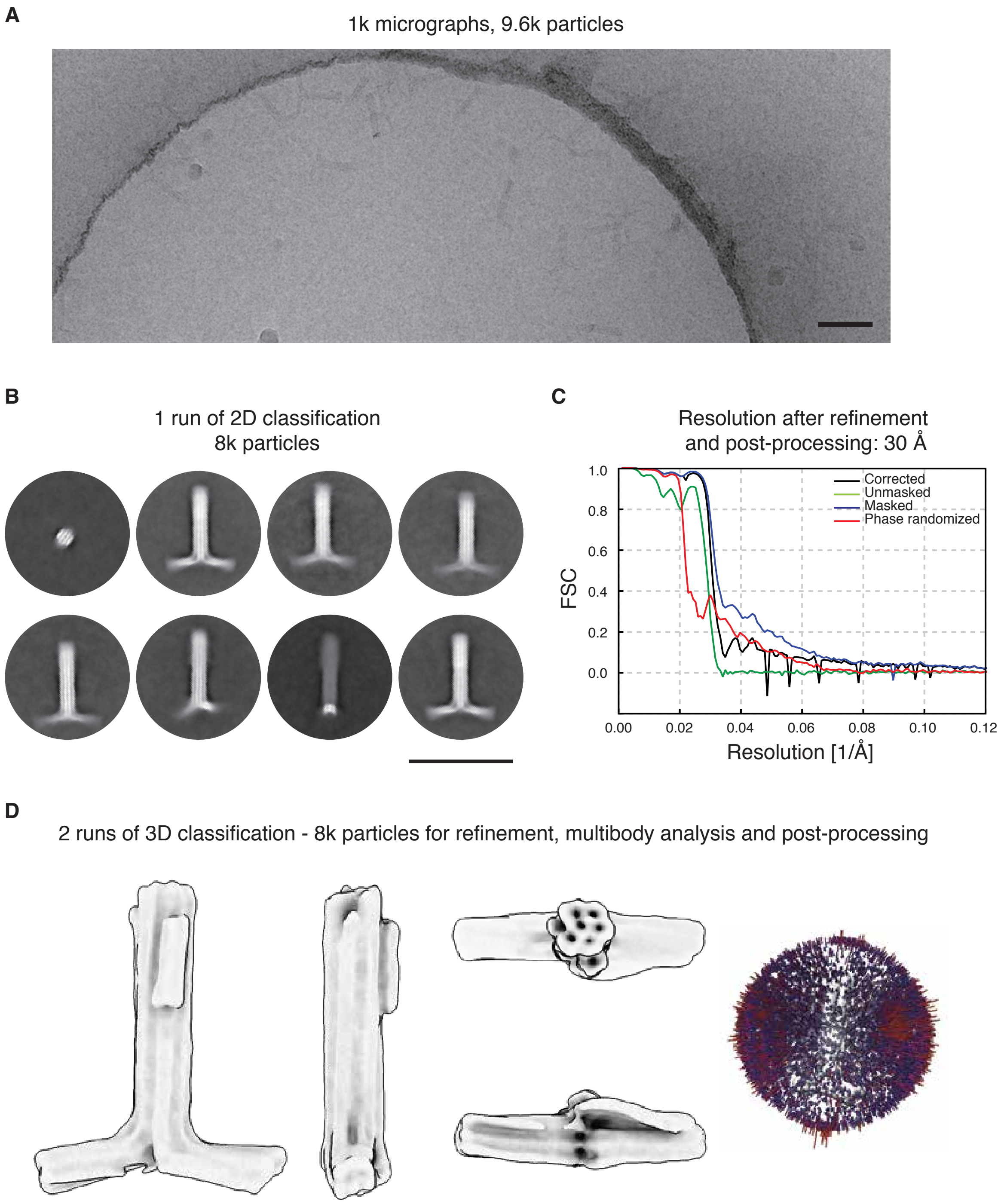


**Figure S10 Cryo EM reconstruction of the camshaft. (A)** Exemplary micrograph. Scale bar 100 nm. **(B)** Exemplary 2D class averages showing the particles in different views. Scale bar 100 nm. **(C)** Fourier Shell Correlation (FSC) of the refined map. **(D)** Post-processed map in different orientations (left) and three-dimensional histogram of the particle orientations (right).

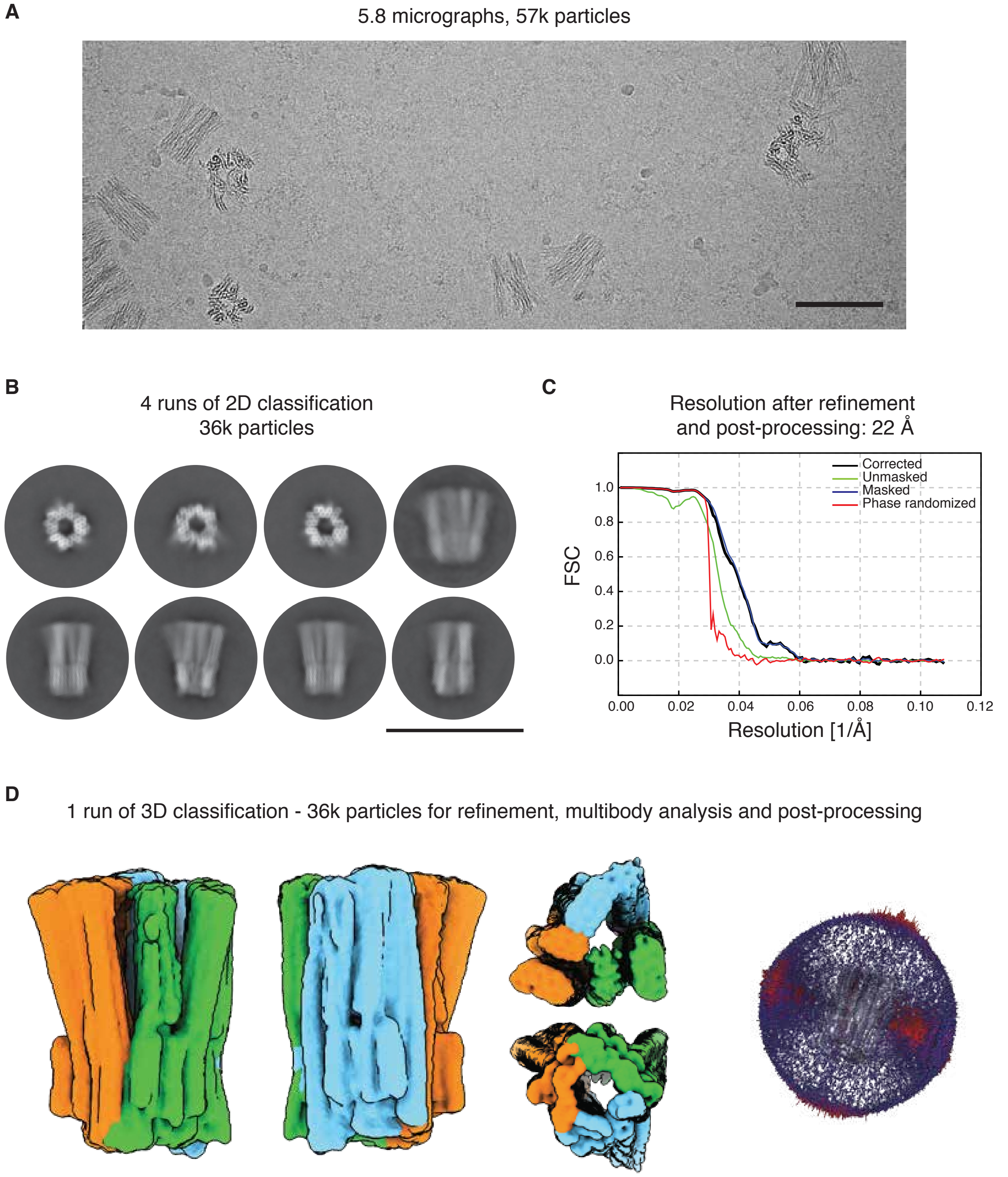


**Figure S11 Cryo EM reconstruction of empty stator. (A)** Exemplary motion-corrected and dose-weighted micrograph. Scale bar 100 nm. **(B)** Exemplary 2D class averages showing the particles in different views. Scale bar 100 nm. **(C)** Fourier Shell Correlation (FSC) of the refined map. **(D)** Composite map from a MultiBody job in different orientations (left) and three-dimensional histogram of the particle orientations (right).

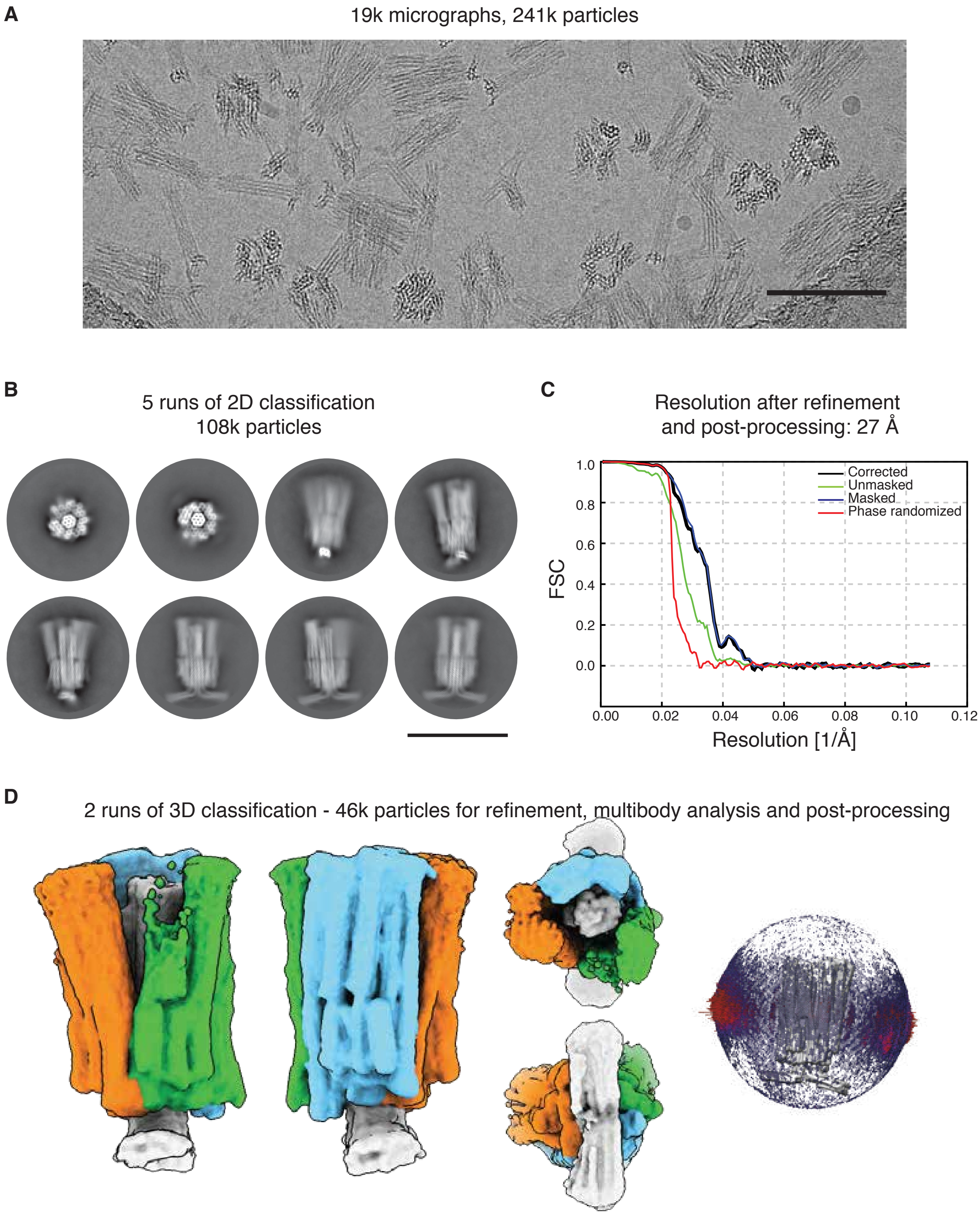


**Figure S12 Cryo EM reconstruction of the rotary mechanism with the camshaft bound to stator unit 1. (A)** Exemplary motion-corrected and dose-weighted micrograph. Scale bar 100 nm. **(B)** Exemplary 2D class averages showing the particles in different views. Scale bar 100 nm. **(C)** Fourier Shell Correlation (FSC) of the refined map. **(D)** Composite map from a MultiBody job in different orientations (left) and three-dimensional histogram of the particle orientations (right).

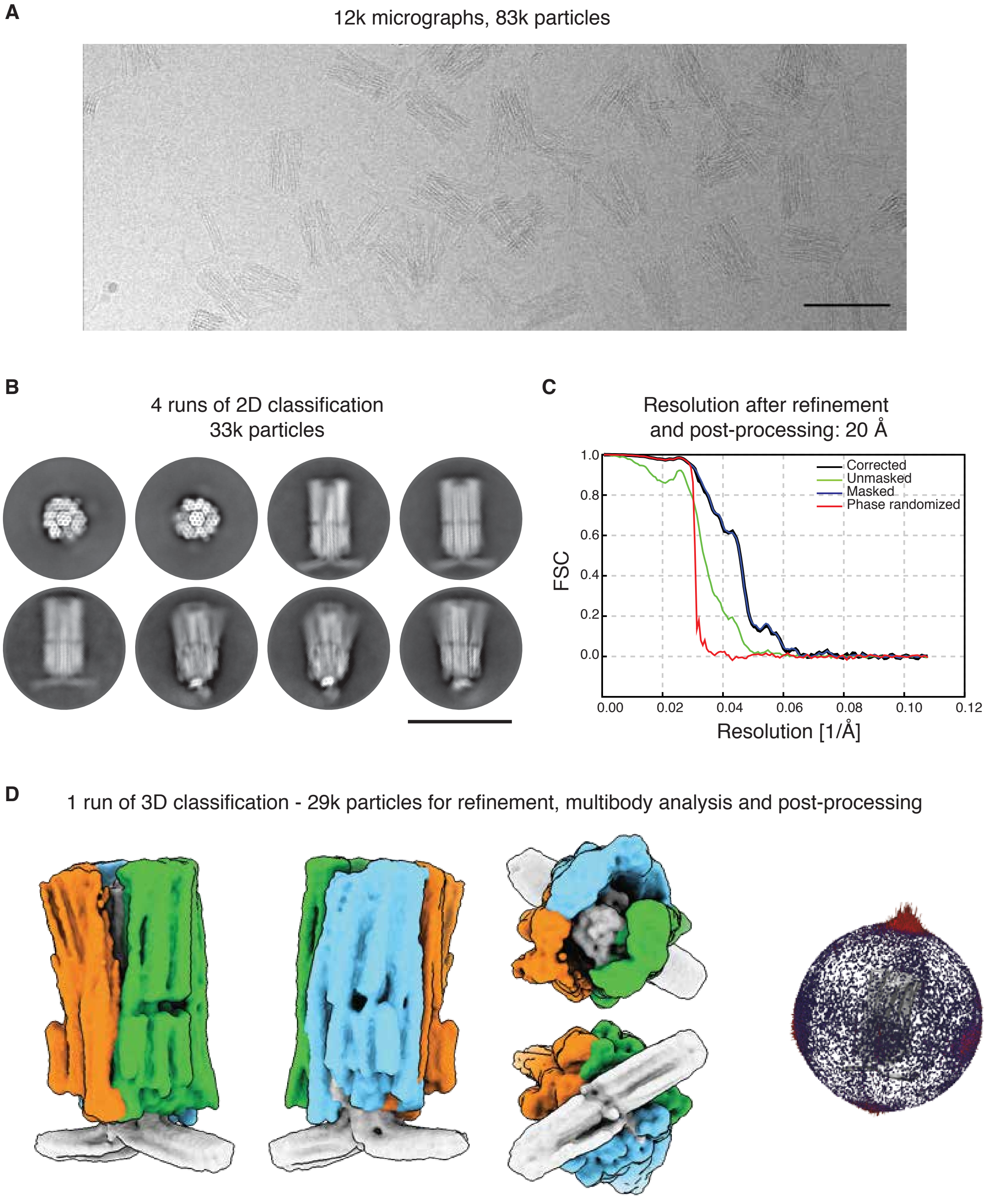


**Figure S13 Cryo EM reconstruction of the rotary complex with the camshaft bound to stator unit 2. (A)** Exemplary motion-corrected and dose-weighted micrograph. Scale bar 100 nm. **(B)** Exemplary 2D class averages showing the particles in different views. Scale bar 100 nm. **(C)** Fourier Shell Correlation (FSC) of the refined map. **(D)** Composite map from a MultiBody job in different orientations (left) and three-dimensional histogram of the particle orientations (right).

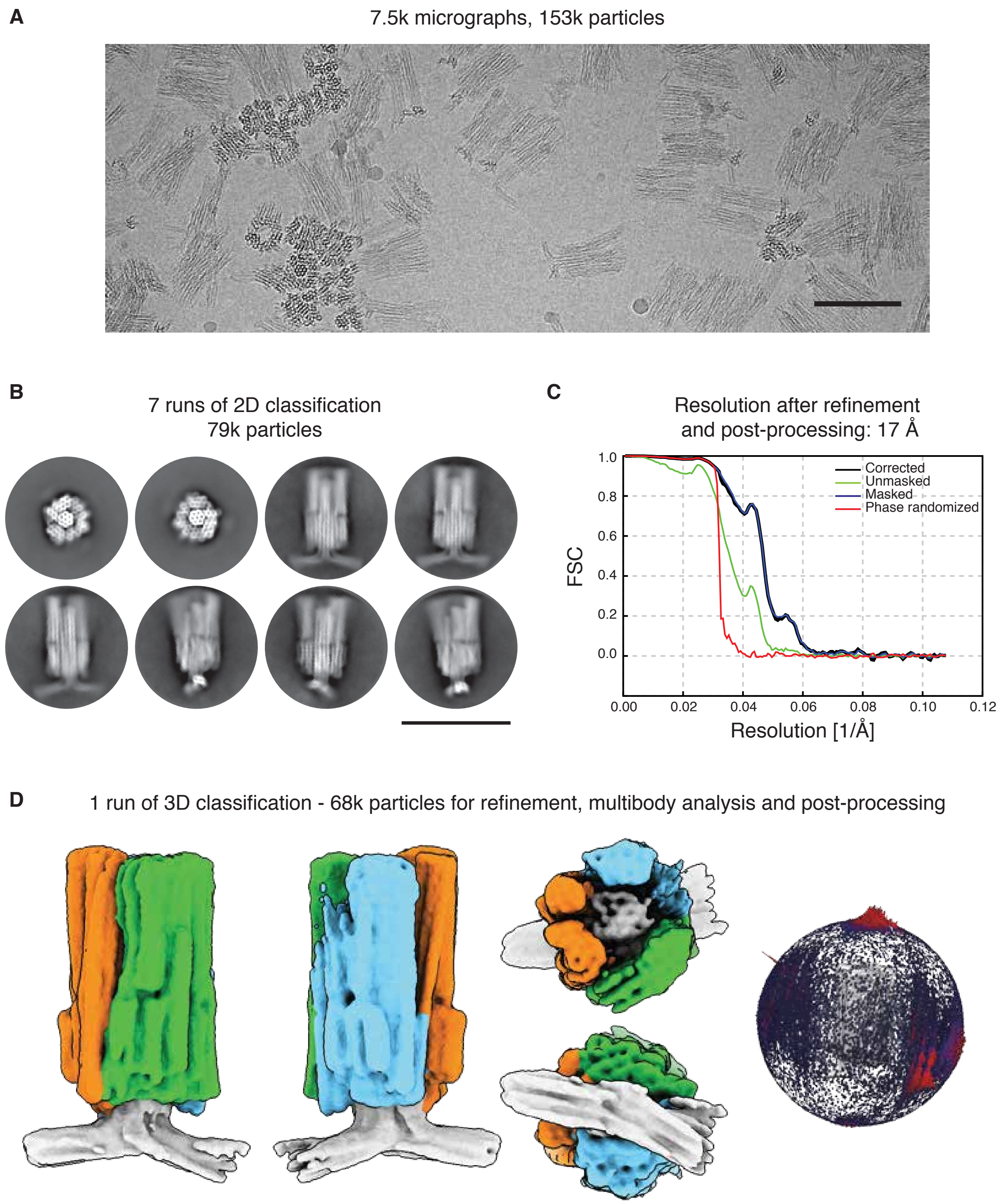


**Figure S14 Cryo EM reconstruction of the rotary complex with the camshaft bound to stator unit 3. (A)** Exemplary motion-corrected and dose-weighted micrograph. Scale bar 100 nm. **(B)** Exemplary 2D class averages showing the particles in different views. Scale bar 100 nm. **(C)** Fourier Shell Correlation (FSC) of the refined map. **(D)** Composite map from a MultiBody job in different orientations (left) and three-dimensional histogram of the particle orientations (right).

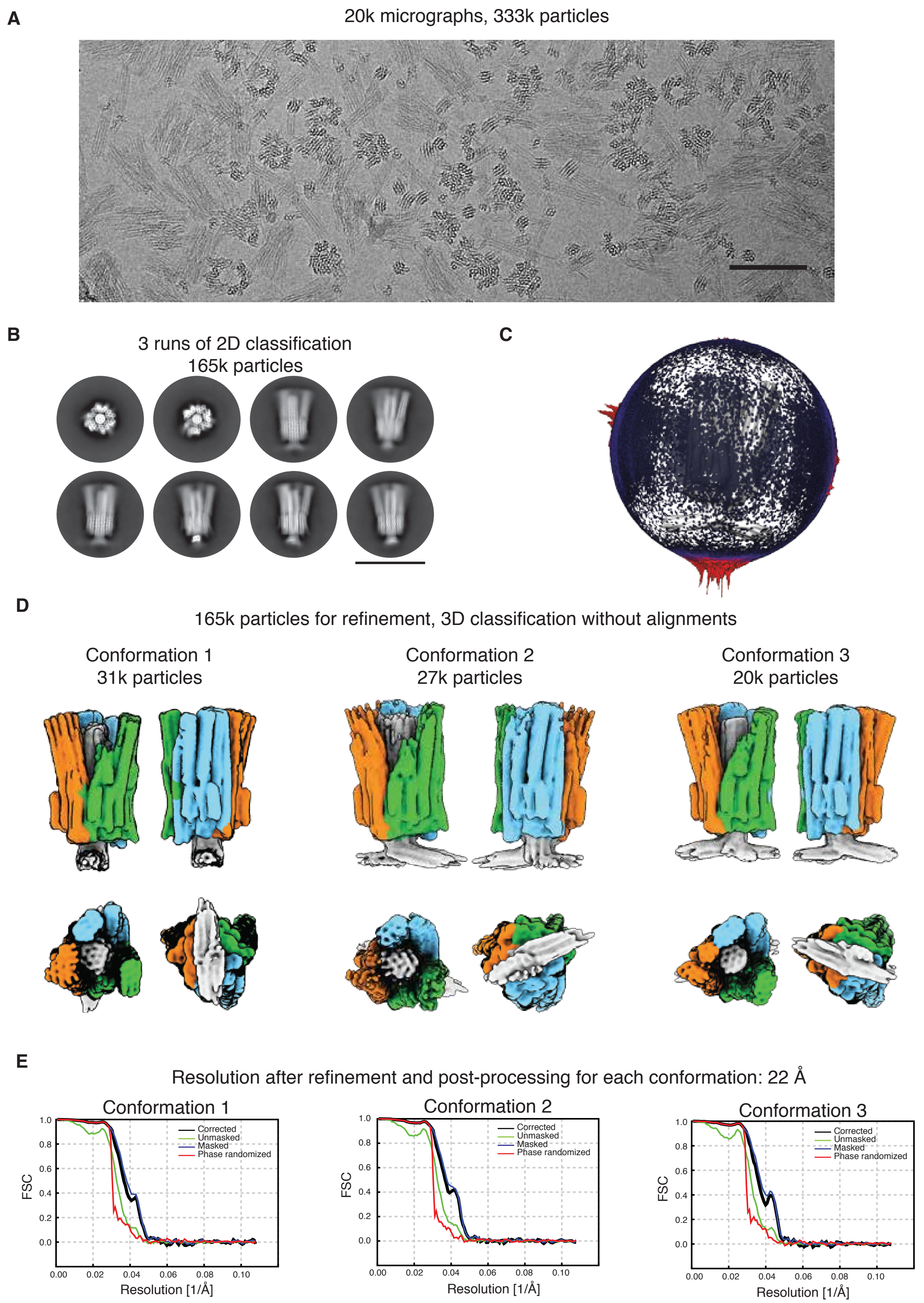


**Figure S15 Cryo EM reconstruction of the rotary complex with the camshaft free to rotate. (A)** Exemplary motion-corrected and dose-weighted micrograph. Scale bar 100 nm. **(B)** Exemplary 2D class averages showing the particles in different views. Scale bar 100 nm. **(C)** Three-dimensional histogram of the particle orientations. **(D)** Composite map from a MultiBody job in different orientations for the three conformations found in the sample. **(E)** Fourier Shell Correlation (FSC) of the refined maps.

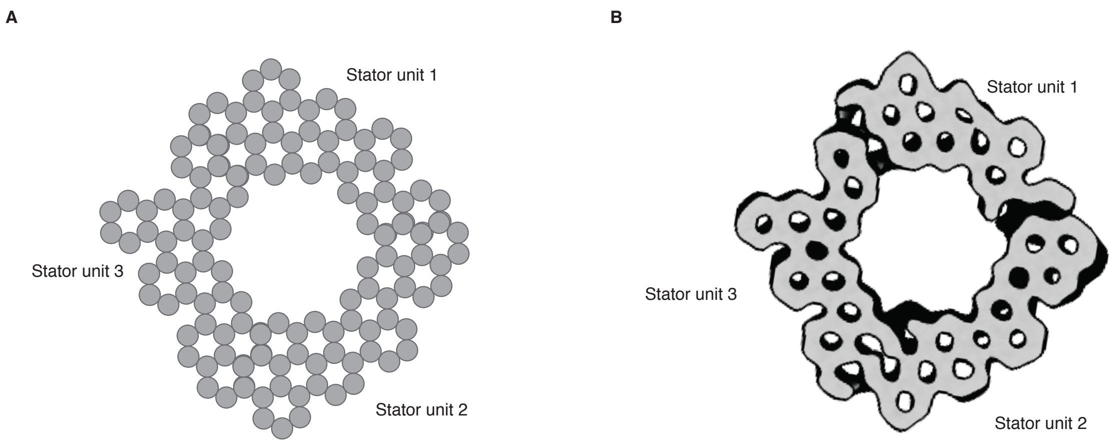


**Figure S16 Stator cross section. (A)** Designed stator top view cross section, where each circle represents a DNA double helix. **(B)** Slice through a 3D reconstruction of the stator (experimental data) showing the stator cross section.

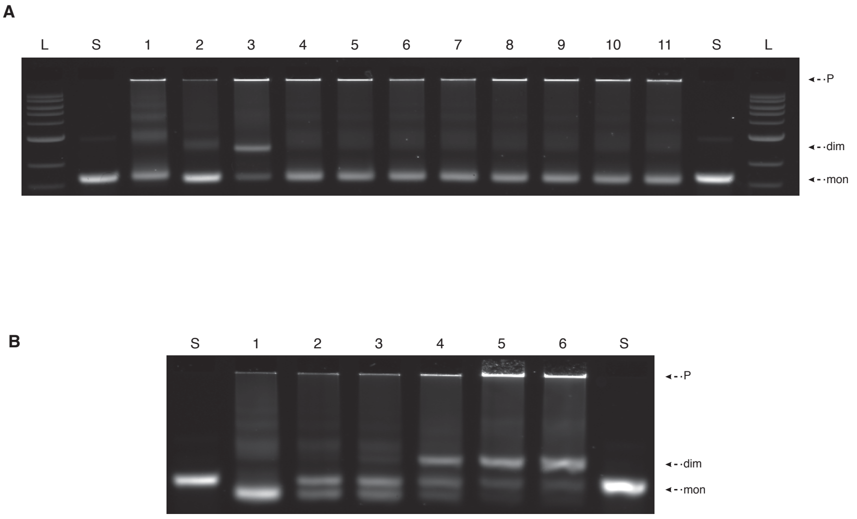


**Figure 17 Camshaft release via toehold mediated strand displacement. (A)** Incubation time screen of invader. Laser-scanned photograph of a 2% agarose gel on which the following samples were electrophoresed: L: ladder; S: p8064 scaffold; lane 1: stator unit 1; lane 2: camshaft; lane 3: stator unit 1 + camshaft dimers; lane 4-11: stator unit 1 + camshaft dimers to which the invader strands were added in 2x excess for different incubation times; lane 4: 0 min; lane 5: 1 min; lane 6: 5 min; lane 7: 10 min; lane 8: 15 min; lane 9: 20 min; lane 10: 25 min; lane 11: 30 min. **(B)** Sub-stoichiometric excess screen of invader strands. Laser-scanned photograph of a 2% agarose gel on which the following samples were electrophoresed: S: p8064 scaffold; lane 1: stator unit 1; lane 2-5: stator unit 1 + camshaft dimers to which the invader strands were added in different excess; lane 2: 2x excess; lane 3: 1x excess; lane 4: 0.4x excess; lane 5: 0.2x excess; lane 6: stator unit 1 + camshaft dimers. P: pockets; dim: dimers; mon: monomers.

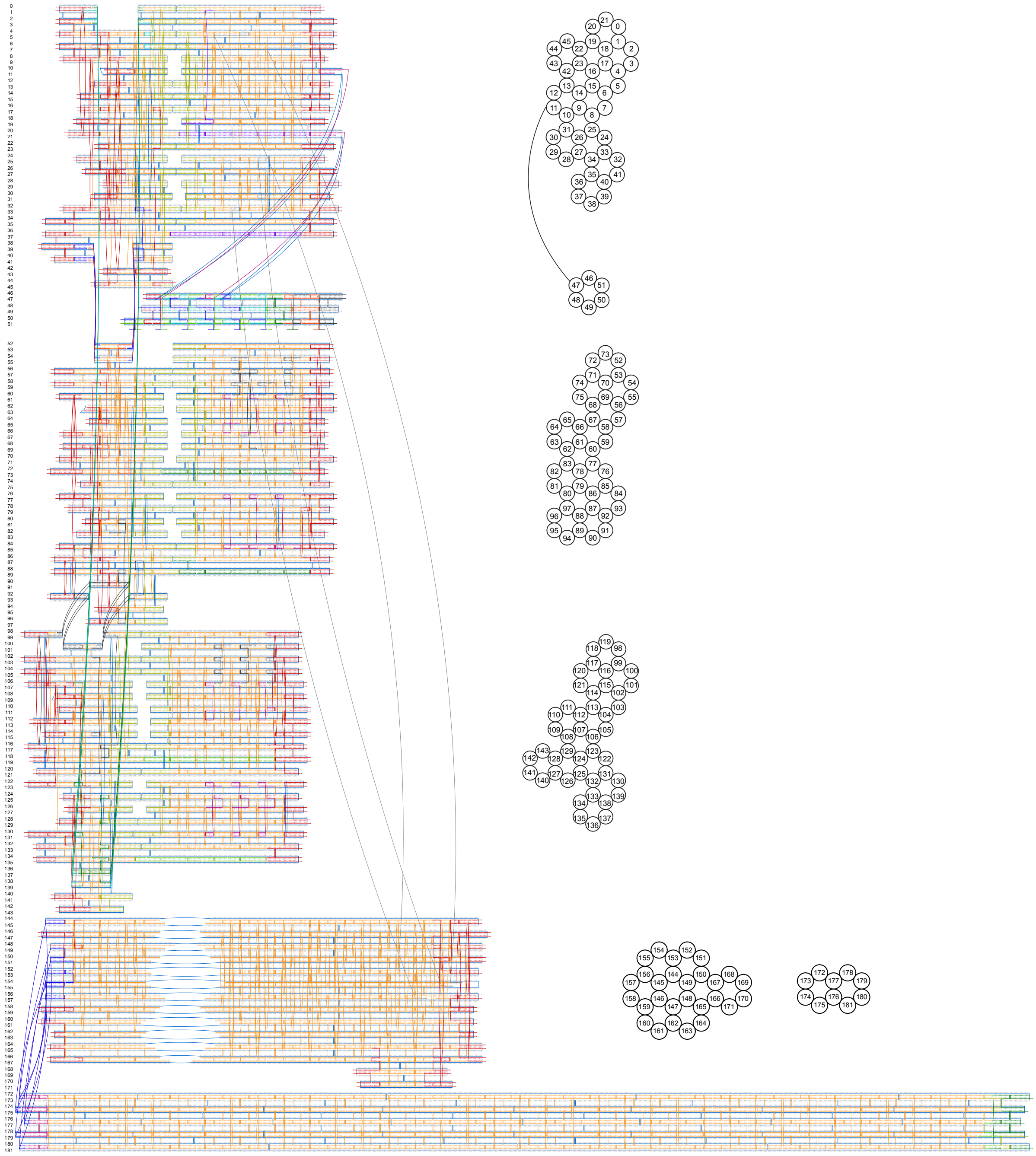

**Figure S18** CaDNAno (*1*) design diagram and bottom view cross sections of the rotary mechanism with modifications for TIRFM analysis.

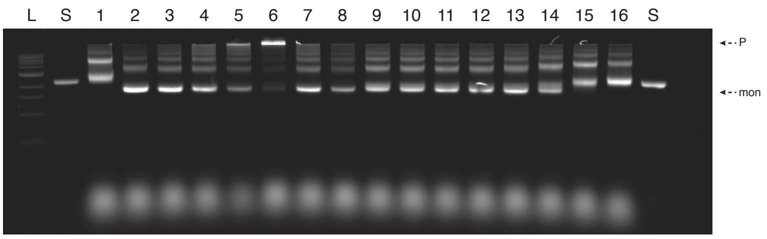


**Figure S19 Folding screen of stator unit 1 with modifications for anchoring to a TIRFM glass slide.** Laser-scanned photograph of a 2% agarose gel on which the following samples were electrophoresed: L: ladder; S: p9072 scaffold; lane 1: 60°C-44°C, 5 mM $MgCl_2$; lane 2: 60°C-44°C, 10 mM $MgCl_2$; lane 3: 60°C-44°C, 15 mM $MgCl_2$; lane 4: 60°C-44°C, 20 mM $MgCl_2$; lane 5: 60°C-44°C, 25 mM $MgCl_2$; lane 6: 60°C-44°C, 30 mM $MgCl_2$; lane 7: 4x staples-to-scaffold excess; lane 8: 10x staples-to-scaffold excess; lane 9: 50°C-47°C, 20 mM $MgCl_2$; lane 10: 52°C-49°C, 20 mM $MgCl_2$; lane 11: 54°C-51°C, 20 mM $MgCl_2$; lane 12: 56°C-53°C, 20 mM $MgCl_2$; lane 13: 58°C-55°C, 20 mM $MgCl_2$; lane 14: 60°C-57°C, 20 mM $MgCl_2$; lane 15: 62°C-59°C; lane 16: 64°C-61°C, 20 mM $MgCl_2$. P: pockets; mon: monomers.

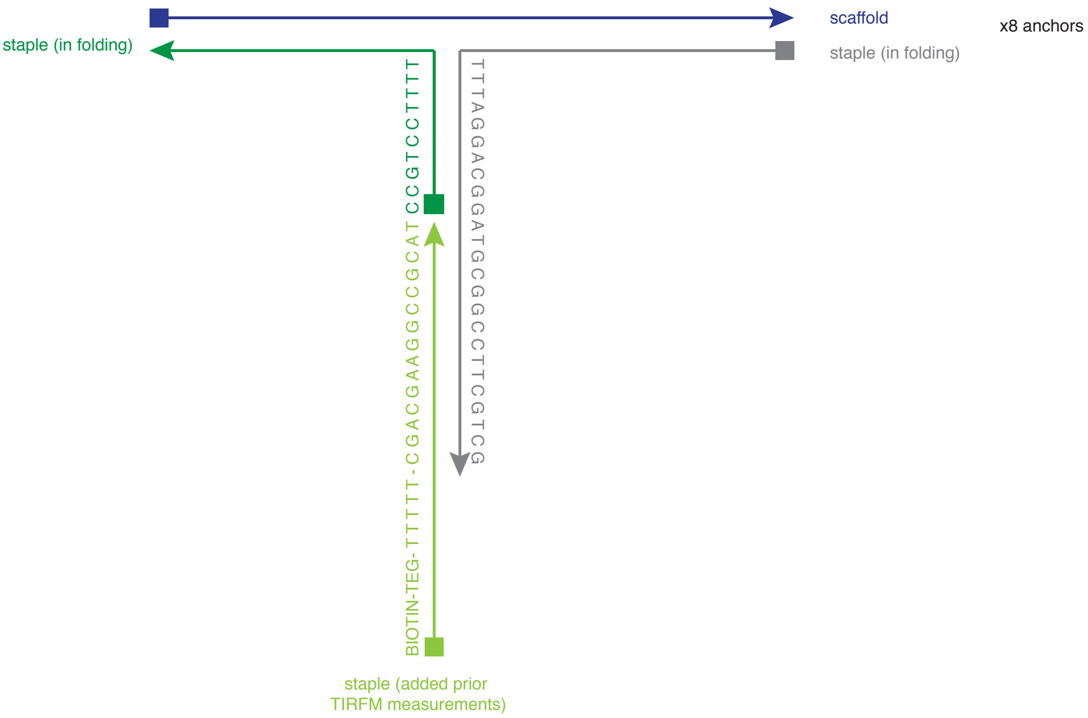


**Figure S20** Schematic representation of the oligos used for anchoring the stator unit 1 on a TIRFM glass slide.

A

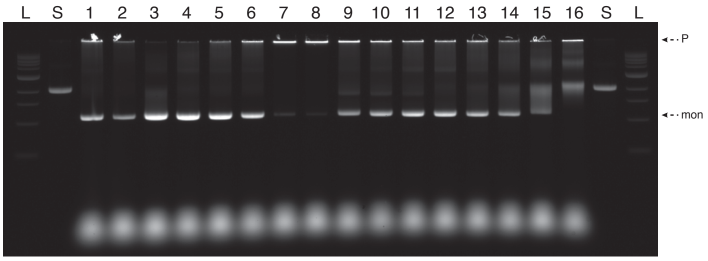


B

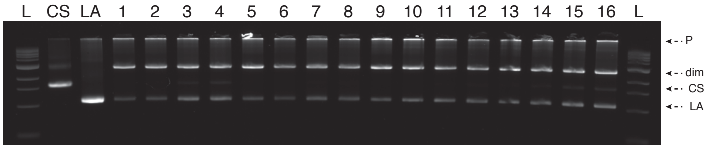


**Figure S21 Folding screen for lever arm and dimerization screen to camshaft. (A)** Folding screen for the lever arm. Laser-scanned photograph of a 2% agarose gel on which the following samples were electrophoresed: L: ladder; S: p8064 scaffold; lane 1: 4x staples-to-scaffold excess; lane 2: 10x staples-to-scaffold excess; lane 3: 60°C-44°C, 5 mM $MgCl_2$; lane 4: 60°C-44°C, 10 mM $MgCl_2$; lane 5: 60°C-44°C, 15 mM $MgCl_2$; lane 6: 60°C-44°C, 20 mM $MgCl_2$; lane 7: 60°C-44°C, 25 mM $MgCl_2$; lane 8: 60°C-44°C, 30 mM $MgCl_2$; lane 9: 50°C-47°C, 20 mM $MgCl_2$; lane 10: 52°C-49°C, 20 mM $MgCl_2$; lane 11: 54°C-51°C, 20 mM $MgCl_2$; lane 12: 56°C-53°C, 20 mM $MgCl_2$; lane 13: 58°C-55°C, 20 mM $MgCl_2$; lane 14: 60°C-57°C, 20 mM $MgCl_2$; lane 15: 62°C-59°C, 20 mM $MgCl_2$; lane 16: 64°C-61°C, 20 mM $MgCl_2$. **(B)** Dimerization screen between the camshaft and the lever arm. Laser-scanned photograph of a 2% agarose gel on which the following samples were electrophoresed: L: ladder; CS: camshaft; LA: lever arm; lane 1: 20 mM $MgCl_2$, RT; lane 2: 30 mM $MgCl_2$, RT; lane 3: 40 mM $MgCl_2$, RT; lane 4: 50 mM $MgCl_2$, RT; lane 5: 20 mM $MgCl_2$, 30°C; lane 6: 30 mM $MgCl_2$, 30°C; lane 7: 40 mM $MgCl_2$, 30°C; lane 8: 50 mM $MgCl_2$, 30°C; lane 9: 20 mM $MgCl_2$, 40°C; lane 10: 30 mM $MgCl_2$, 40°C; lane 11: 40 mM $MgCl_2$, 40°C; lane 12: 50 mM $MgCl_2$, 40°C; lane 13: 20 mM $MgCl_2$, 50°C; lane 14: 30 mM $MgCl_2$, 50°C; lane 15: 40 mM $MgCl_2$, 50°C; lane 16: 50 mM $MgCl_2$, 50°C. P: pockets; dim: dimers; mon: monomers.

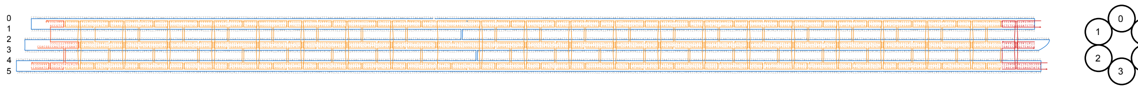


**Figure S22** CaDNAno (*1*) design diagram (left) and bottom-view cross section (right) of the prolongation of the 6hb in stator unit 1.

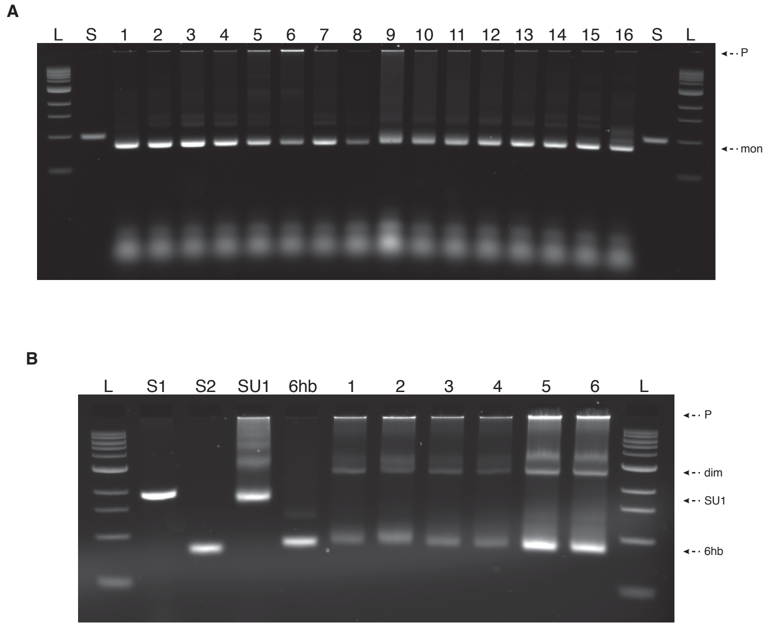


**Figure S23 Folding screen for the 6hb and dimerization screen with stator unit 1. (A)** Folding screen for the 6hb. Laser-scanned photograph of a 2% agarose gel on which the following samples were electrophoresed: L: ladder; S: p2873 scaffold; lane 1: 60°C-44°C, 5 mM $MgCl_2$; lane 2: 60°C-44°C, 10 mM $MgCl_2$; lane 3: 60°C-44°C, 15 mM $MgCl_2$; lane 4: 60°C-44°C, 20 mM $MgCl_2$; lane 5: 60°C-44°C, 25 mM $MgCl_2$; lane 6: 60°C-44°C, 30 mM $MgCl_2$; lane 7: 4x staples-to-scaffold excess; lane 8: 10x staples-to-scaffold excess; lane 9: 50°C-47°C, 20 mM $MgCl_2$; lane 10: 52°C-49°C, 20 mM $MgCl_2$; lane 11: 54°C-51°C, 20 mM $MgCl_2$; lane 12: 56°C-53°C, 20 mM $MgCl_2$; lane 13: 58°C-55°C, 20 mM $MgCl_2$; lane 14: 60°C-57°C, 20 mM $MgCl_2$; lane 15: 62°C-59°C; lane 16: 64°C-61°C, 20 mM $MgCl_2$. **(B)** Dimerization screen between the stator unit 1 and the 6hb. Laser-scanned photograph of a 2% agarose gel on which the following samples were electrophoresed: L: ladder; S1: p9072 scaffold; S2: p2873 scaffold; SU1: stator unit 1; 6hb: 6 helix-bundle; lane 1: 40 mM $MgCl_2$, 30°C; lane 2: 50 mM $MgCl_2$, 30°C; lane 3: 40 mM $MgCl_2$, 40°C; lane 4: 50 mM $MgCl_2$, 40°C; lane 5: 40 mM $MgCl_2$, 50°C; lane 6: 50 mM $MgCl_2$, 50°C. P: pockets; mon: monomers; dim: dimers.

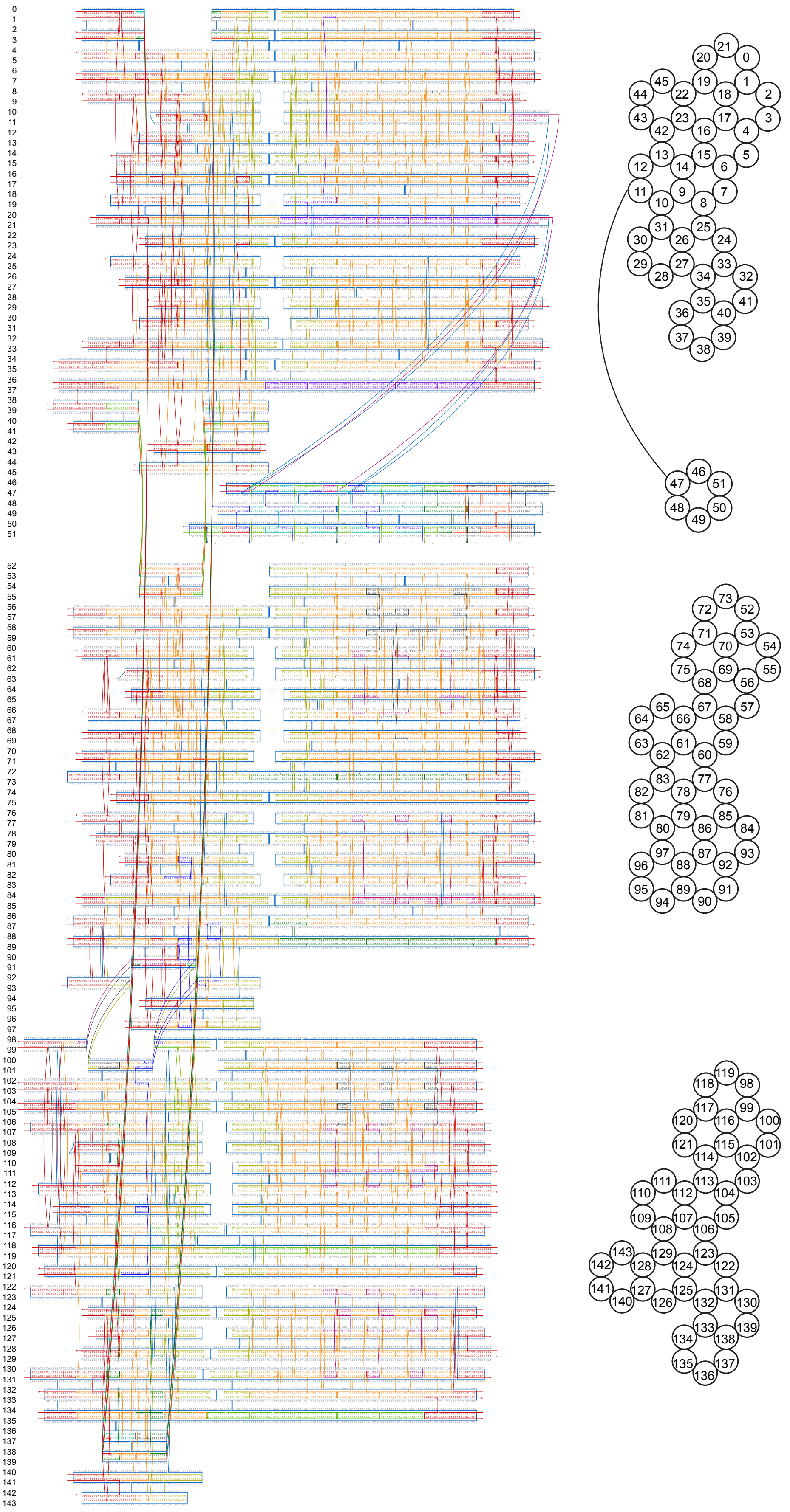

**Figure S24** CaDNAno (*1*) design diagram (left) and bottom view cross sections (right) of v2 stator units.

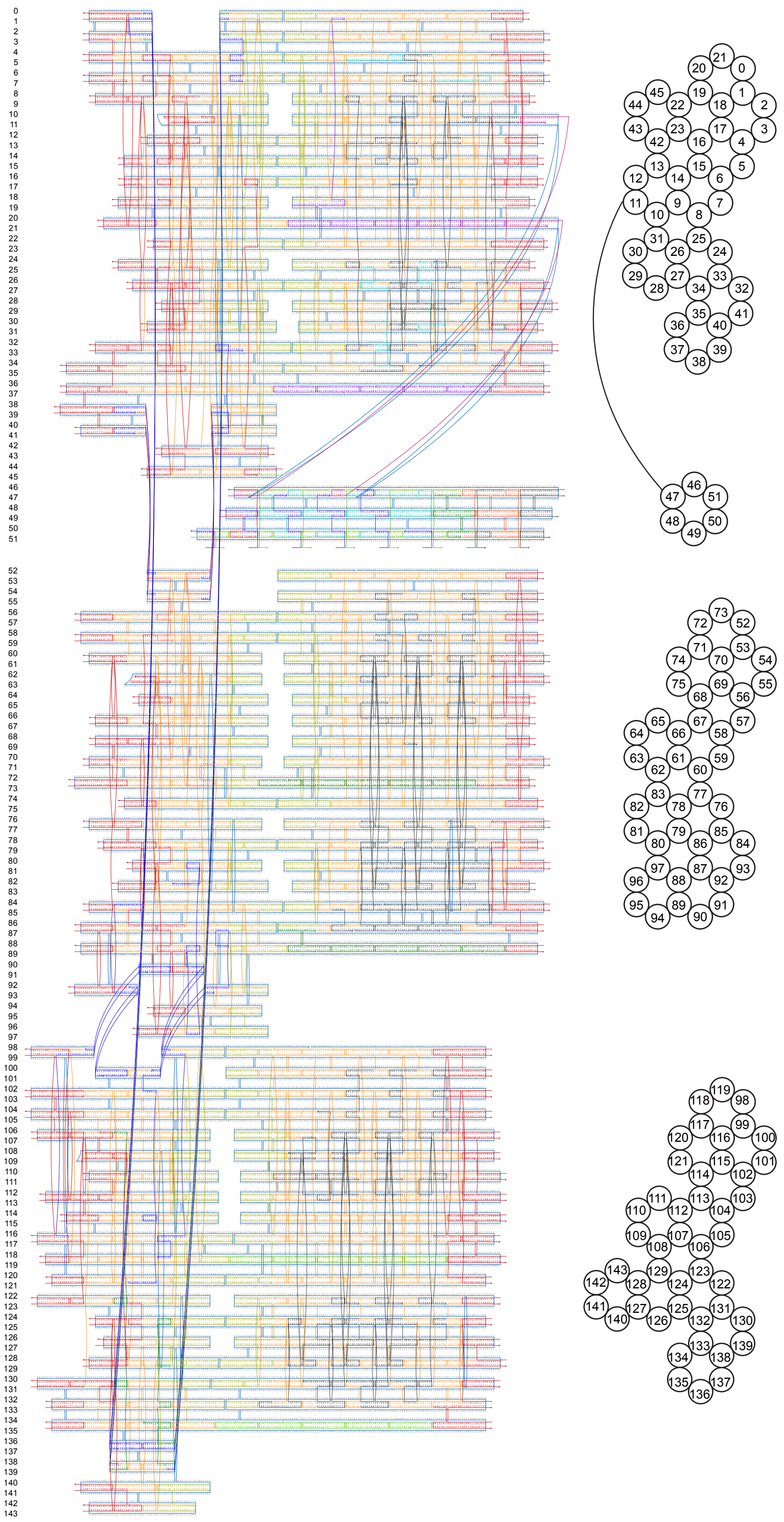


**Figure S25** CaDNAno (*1*) design diagram (left) and bottom view cross sections (right) of v3 stator units.

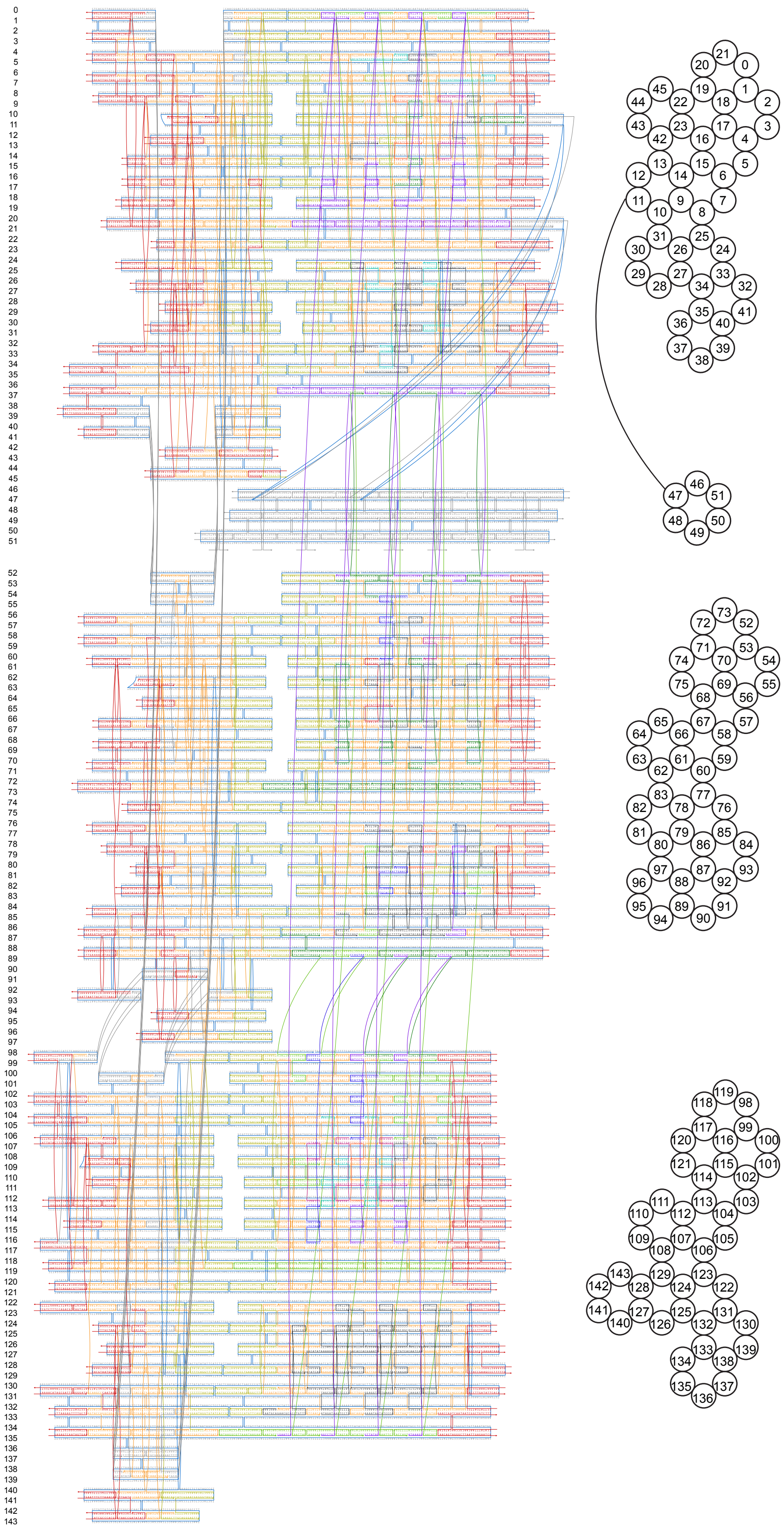

**Figure S26** CaDNAno (*1*) design diagram (left) and bottom view cross sections (right) of v4 stator units.

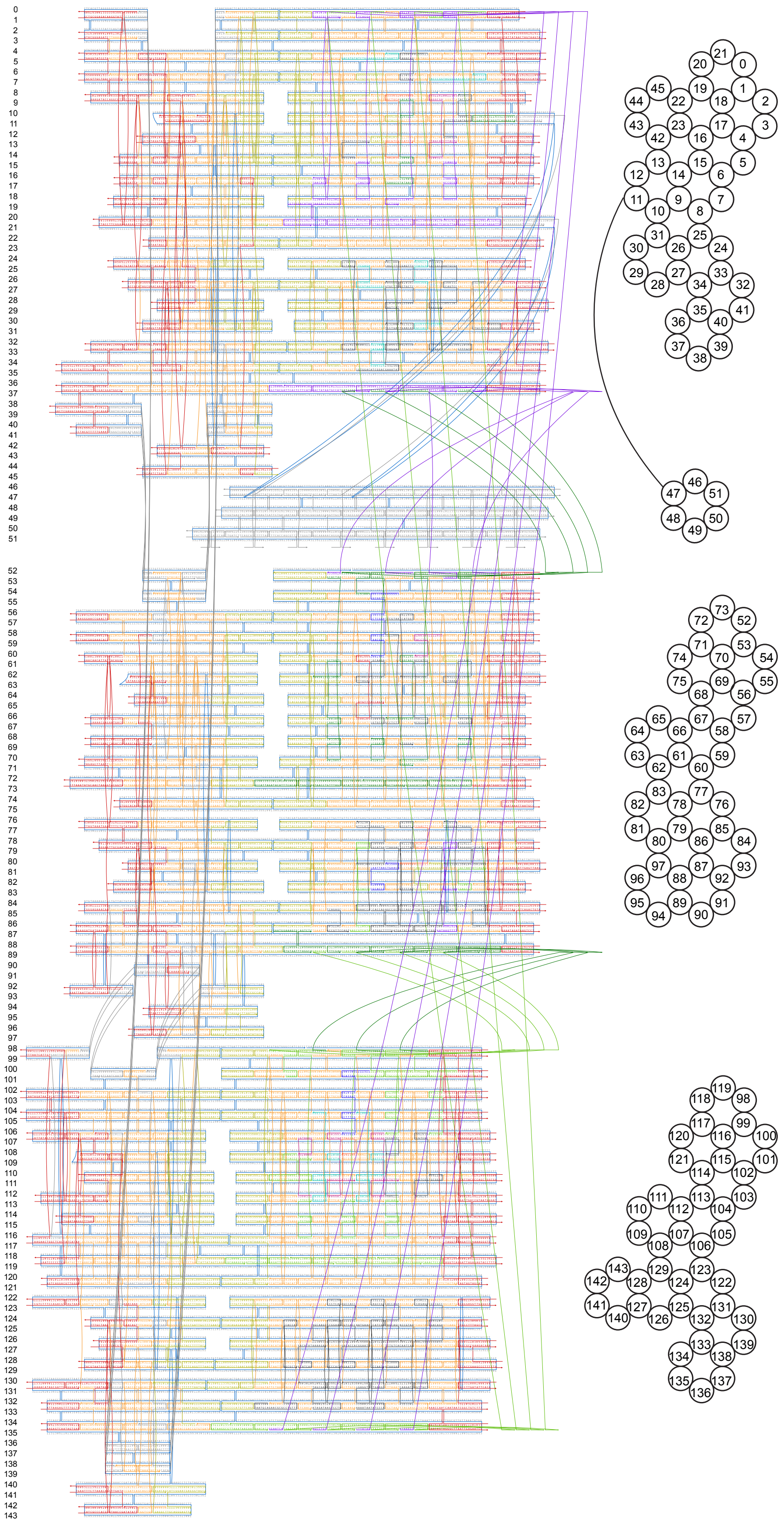

**Figure S27** CaDNAno (*1*) design diagram (left) and bottom view cross sections (right) of v5 stator units. The spacer oligos between the stator units are 25 Ts long.

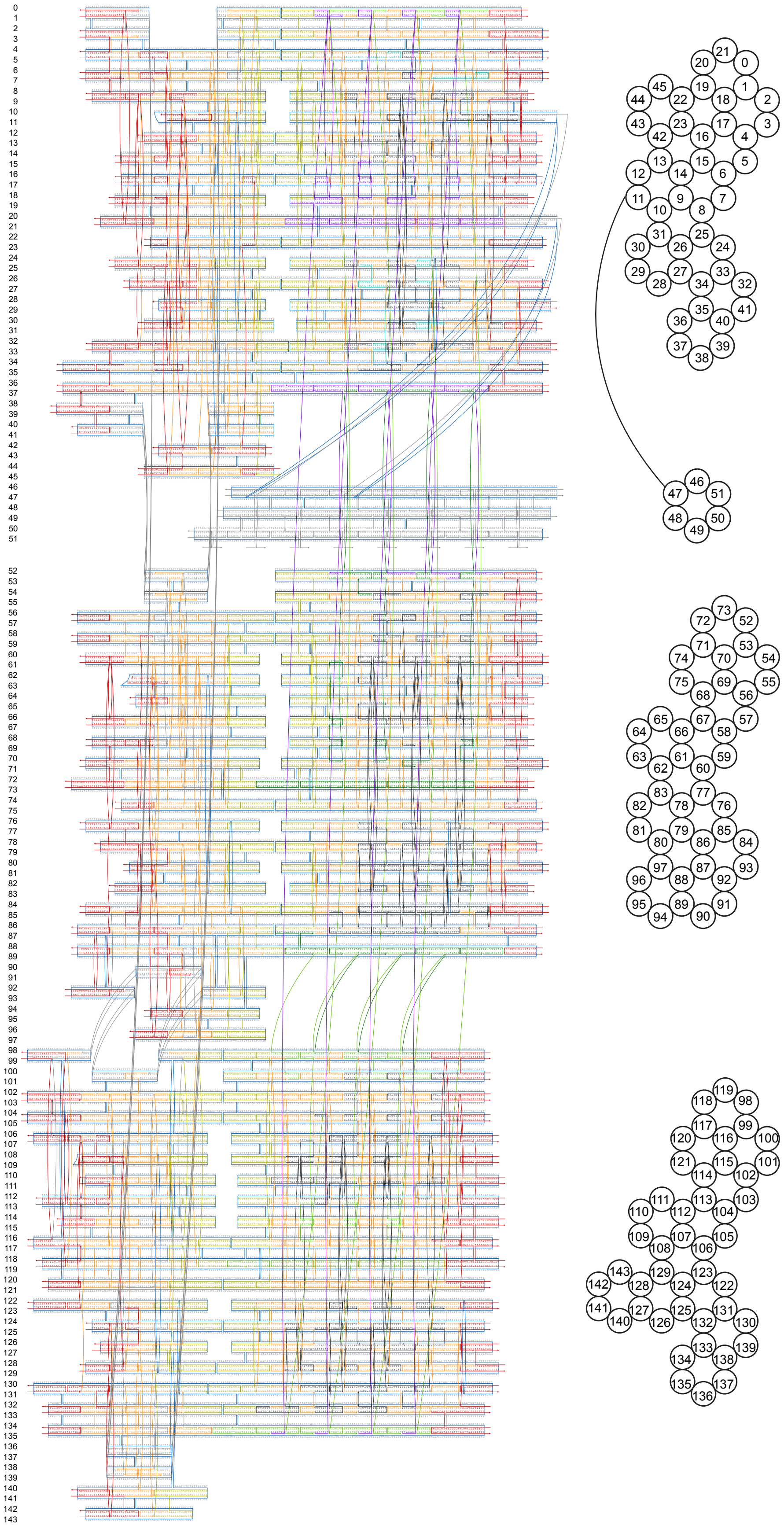

**Figure S28** CaDNAno (*1*) design diagram (left) and bottom view cross sections (right) of v6 stator units.

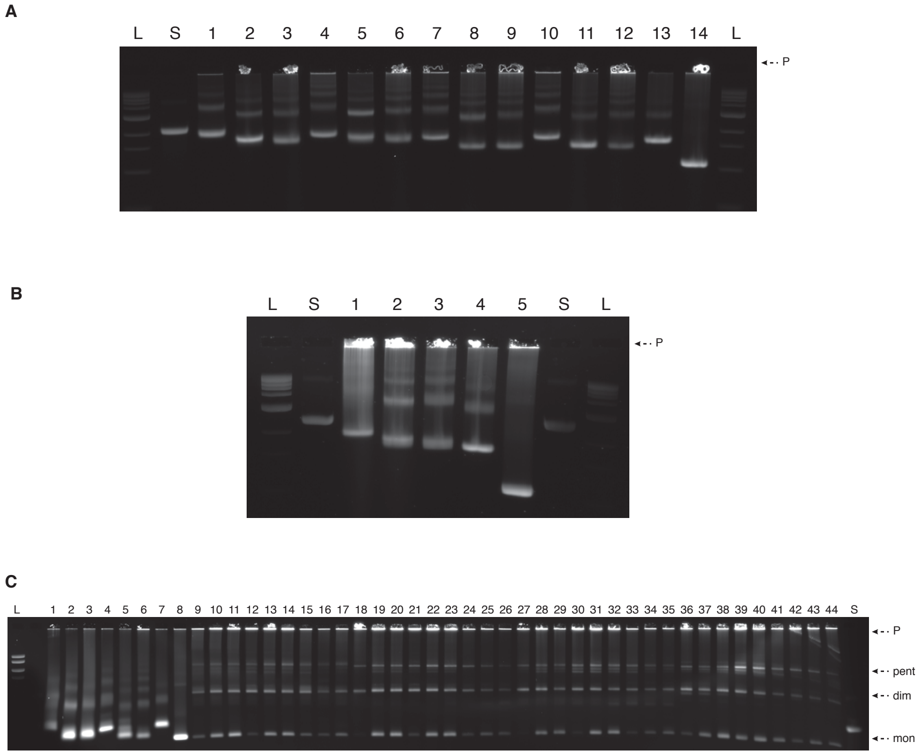


**Figure S29 Folding and polymerization of the structure modified to have a stiffer or a more flexible stator. (A)** Folding of the different monomers. Laser-scanned photograph of a 2% agarose gel on which the following samples were electrophoresed: L: ladder; S: p8064 scaffold; lane 1: stator unit 1, v2; lane 2: stator unit 2, v2; lane 3: stator unit 3, v2; lane 4: stator unit 1, v4; lane 5: stator unit 2, v4; lane 6: stator unit 3, v4; lane 7: stator unit 1, v3; lane 8: stator unit 2, v3; lane 9: stator unit 3, v3; lane 10: stator unit 1, v6; lane 11: stator unit 2, v6; lane 12: stator unit 3, v6; lane 13: camshaft; lane 14: lever arm. **(B)** Folding of v5. Laser-scanned photograph of a 2% agarose gel on which the following samples were electrophoresed: L: ladder; S: p8064 scaffold; lane 1: stator unit 1; lane 2: stator unit 2; lane 3: stator unit 3; lane 4: camshaft; lane 5: lever arm. **(C)** Polymerization screen of the modified stator variants. Laser-scanned photograph of a 2% agarose gel on which the following samples were electrophoresed: L: ladder; lane 1: stator unit 1, v2; lane 2: stator unit 2, v2; lane 3: stator unit 3, v2; lane 4: stator unit 1, v6; lane 5: stator unit 2, v6; lane 6: stator unit 3, v6; lane 7: camshaft; lane 8: lever arm; lane 9: v2, 30 mM $MgCl_2$, 30°C; lane 10: v2, 40 mM $MgCl_2$, 30°C; lane 11: v2, 50 mM $MgCl_2$, 30°C; lane 12: v2, 30 mM $MgCl_2$, 40°C; lane 13: v2, 40 mM $MgCl_2$, 40°C; lane 14: v2, 50 mM $MgCl_2$, 40°C; lane 15: v2, 30 mM $MgCl_2$, 50°C; lane 16: v2, 40 mM $MgCl_2$, 50°C; lane 17: v2, 50 mM $MgCl_2$, 50°C; lane 18: v4, 30 mM $MgCl_2$, 30°C; lane 19: v4, 40 mM $MgCl_2$, 30°C; lane 20: v4, 50 mM $MgCl_2$, 30°C; lane 21: v4, 30 mM $MgCl_2$, 40°C; lane 22: v4, 40 mM $MgCl_2$, 40°C; lane 23: v4, 50 mM $MgCl_2$, 40°C; lane 24: v4, 30 mM $MgCl_2$, 50°C; lane 25: v4, 40 mM $MgCl_2$, 50°C; lane 26: v4, 50 mM $MgCl_2$, 50°C; lane 27: v3, 30 mM $MgCl_2$, 30°C; lane 28: v3, 40 mM $MgCl_2$, 30°C; lane 29: v3, 50 mM $MgCl_2$, 30°C; lane 30: v3, 30 mM $MgCl_2$, 40°C; lane 31: v3, 40 mM $MgCl_2$, 40°C; lane 32: v3, 50 mM $MgCl_2$, 40°C; lane 33: v3, 30 mM $MgCl_2$, 50°C; lane 34: v3, 40 mM $MgCl_2$, 50°C; lane 35: v3, 50 mM $MgCl_2$, 50°C; lane 36: v6, 30 mM $MgCl_2$, 30°C; lane 37: v6, 40 mM $MgCl_2$, 30°C; lane 38: v6, 50 mM $MgCl_2$, 30°C; lane 39: v6, 30 mM $MgCl_2$, 40°C; lane 40: v6, 40 mM $MgCl_2$, 40°C; lane 41: v6, 50 mM $MgCl_2$, 40°C; lane 42: v6, 30 mM $MgCl_2$, 50°C; lane 43: v6, 40 mM $MgCl_2$, 50°C; lane 44: v6, 50 mM $MgCl_2$, 50°C. P: pockets; pent: pentamers; dim: dimers; mon: monomers.

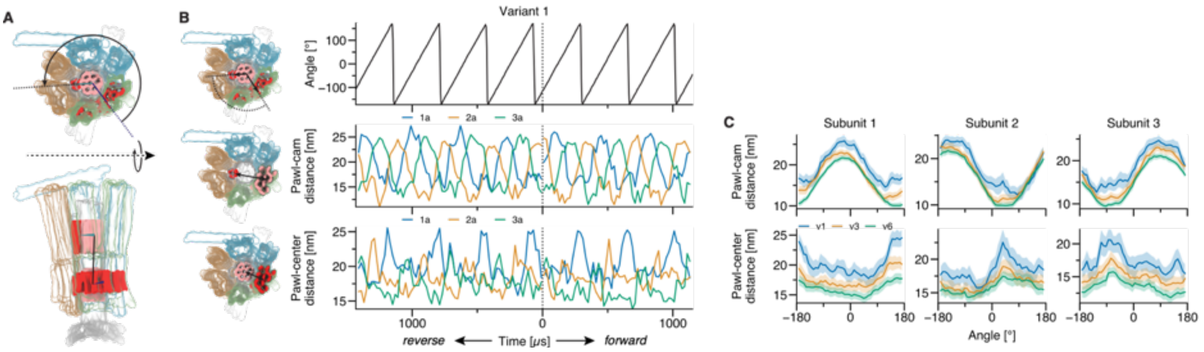


**Figure S30 Simulation of driven rotation. (A)** Schematic of the dihedral angle used to drive the rotation. To drive the rotation, the rest angle of a harmonic potential applied to this dihedral angle was increased or decreased with a constant rate. **(B)** Forced rotation of variant 1. The time-varying potential acting on the dihedral angle described in panel A caused the rotor to spin (top). The cam cyclically approached each pawl (middle), causing it to deform away from the center of the rotor (bottom). Schematics of each metric are shown to the left of the plots. **(C)** Comparison of forced rotation of variants 1, 3 and 6.

### Supplementary Movies

**Supplementary Movie 1:** Coarse-grained mrDNA simulations of all six rotor variants with a ~5 bp/bead resolution model. Starting from an idealized geometry taken from the CaDNAno (*1*) design, each simulation lasted 20 µs.

**Supplementary Movie 2:** Coarse-grained mrDNA simulations of driven rotation of variants 1, 3 and 6 with a ~5 bp/bead resolution model. Starting from an equilibrated configuration, the rotation angle was increased or decreased at a rate of 1 degree per 20 µs. Simulations lasted at least 20 ms, or approximately 3 rotations in either direction.

**Supplementary Movie 3:** Average configuration of the rotor variants 1, 2 and 3 at a given rotation angle. The coordinates of the rotor were first aligned to minimize the root mean square deviation of the bearing region below the pawls, and were subsequently sorted according to the instantaneous value of the rotation angle into 10° bins. The ensemble of configurations within each bin was averaged and is depicted in the animation. The average includes configuration sampled from both forward and reverse driven rotation simulations.
